# Prefrontal cortex encodes behavior states decoupled from movement

**DOI:** 10.1101/2024.11.11.623038

**Authors:** Ida Välikangas Rautio, Fredrik Nevjen, Ingeborg Hem, Benjamin A. Dunn, Jonathan R. Whitlock

## Abstract

Prefrontal cortex is often viewed as an extension of the motor system, but little is understood of how it relates to natural motor behavior. We therefore tracked the kinematics of freely moving rats performing minimally structured tasks and measured which aspects of behavior were read out in prefrontal neural populations. Naturalistic behaviors such as rearing or chasing a bait were each encoded by unique neural ensembles, but the behavioral representations were not anchored to posture or movement. Rather, the coding of kinematic features depended on their relevance to the animals’ current behavior or which task the animal performed. Behavior-specific ensembles often preceded and outlasted physical actions and, accordingly, prefrontal population activity evolved at slower timescales than in motor cortex. These findings argue that prefrontal coding of behavior is not locked to motor output, and may instead reflect motivations to perform certain actions rather than the actions themselves.

**Highlights:**

- Prefrontal neural ensembles uniquely encode different naturalistic actions
- Behavioral tuning is not explained by movement kinematics
- Population activity in prefrontal cortex evolves slower than in M1
- Single-cell coding of behavior varies across tasks yet ensemble coding is stable

## Introduction

The prefrontal cortices sit atop an executive hierarchy that ultimately generates motor behavior to meet specific goals (Prescott et al., 1999; Fuster, 2001; Botvinick et al., 2009; Fuster, 2015; Fine & Hayden, 2021). On this view, prefrontal regions at the upper end of the hierarchy represent internal schema that structure action selection, and subsumed motor pathways issue commands to enact actions. However, despite its central role in behavioral control, little is understood of how prefrontal activity relates to the pacing, structure or content of self-generated naturalistic actions.

One reason for this is that prefrontal cortex is typically studied in trial-based tasks where restrained subjects perform simple movements to report cognitive outcomes (Gomez-Marin et al., 2014; Nastase et al., 2020). Such approaches are effective at isolating cognitive functions but constrain the dimensionality and timing of behavior, which eclipses the bigger picture of how prefrontal activity relates to behavior at different timescales. More fundamentally, movement drives brain-wide changes in neural activity often larger than those driven by task-related features (Stringer et al., 2019; Mussall et al., 2019; Steinmetz et al., 2019; Aimon et al., 2019; Salkoff et al., 2020; Chen et al, 2024), which complicates disentangling of cognitive and motor signals once a subject is moving.

Defining how prefrontal activity relates to motor behavior could therefore benefit from studying subjects outside of tasks that pair cognition with pre- determined actions. Recording from freely behaving animals, for example, could be effective, since the irregularity of unscripted movements affords a more transparent readout of kinematic effects on neural activity (Tremblay et al., 2023; Voloh et al., 2023). The lack of a temporally rigid task may also unmask differences in coding timescales for low-level motor features and cognitive processes, as found for several types of sensory processing (Yang & Zador, 2012; Honey et al., 2012; Papo, 2013; Murray et al., 2014; Runyan & Harvey, 2017; Purandare & Mehta, 2023), but not yet studied for naturalistic motor behavior. Using freely moving subjects also permits the analysis of prefrontal activity during more complex behaviors (Reinhold et al., 2019; Bagi et al., 2022; Tremblay et al., 2023; Voloh et al., 2023), which sheds more light on the role of prefrontal cortex in animals’ behavioral ecology.

We therefore performed Neuropixels recordings in medial prefrontal cortex (mPFC) in unrestrained rats, and tracked them in 3D while they engaged in three minimally structured tasks. We used unbiased analyses of population coactivity to identify salient behavioral events, which revealed that species-typical actions, such as active investigation, rearing, or chasing after food (Karli, 1956; Timberlake, 1983; Renner & Seltzer, 1991; Thinus-Blanc, 1996; Lever et al., 2006; Sturman et al., 2018; Layfield et al., 2023; Shan et al., 2023) were each associated with their own patterns of ensemble activity. We identified seven such ”behavior states” that could be decoded from population activity in each animal, with each state defined by a unique combination of kinematic features. However, neural tuning to those features was not hard-coded, but gated by whichever momentary behavior the animal performed. For example, the neural coding of speed was stronger when animals engaged in bait-chasing, but less than half the ”chasing” neurons stably encoded speed *per se* outside of chasing. Moreover, neural ensembles often formed before or between bouts of their preferred behavior, such as rearing, and population activity in mPFC evolved more slowly than in M1. Single-cell behavioral selectivity also varied across different tasks, suggesting stronger influences of local affordances rather than movement kinematics. We also uncovered head-direction tuning in mPFC that, in contrast with behavior state representations, appeared stable across tasks and behaviors. Thus, whereas the rest of sensory and motor cortices are tightly linked to the physical expression of motor output (Musall et al., 2019; Stringer et al., 2019; Salkoff et al., 2020; Mimica et al., 2023), prefrontal areas are not, and might instead reflect cognitive or motivational states that drive actions depending on the task at hand.

## Results

### Fluctuations in prefrontal population activity signify naturalistic behaviors

We recorded neural activity in mPFC with chronically implanted Neuropixels 1.0 probes while freely moving adult female rats (n = 3) engaged in three different behavioral tasks. The animals were habituated to the recording environment but had minimal prior training, and were tracked in 3D via retroreflective markers on the back and head (Methods; Mimica et al., 2018). In the first task, animals freely explored a 1.9 m long elevated track with 11 cm high walls and a solid mesh divider in the middle, with a familiar female rat on the other side and tracked the same way (Figure 1A, left; Video S1). The divider allowed the animals to interact socially but not climb on each other; they were otherwise free to explore the track and climb the track walls. The second task was foraging in a 1.25 x 1.9 m open arena with 50 cm high walls (Figure 1A, left; Video S1). Third was a chasing task in the same arena, which included bouts when animals chased a bait (puffed corn with a retroreflective marker above it) suspended from a small fishing rod and continuously moved semi-randomly by the experimenter (Video S1). Without training, the animals actively pursued the bait for as long as it was in the arena, and chasing bouts (4-15 per session) were separated by quiescent periods when the bait was removed and animals consumed the reward (Figure 1A, left).

**Figure 1.**
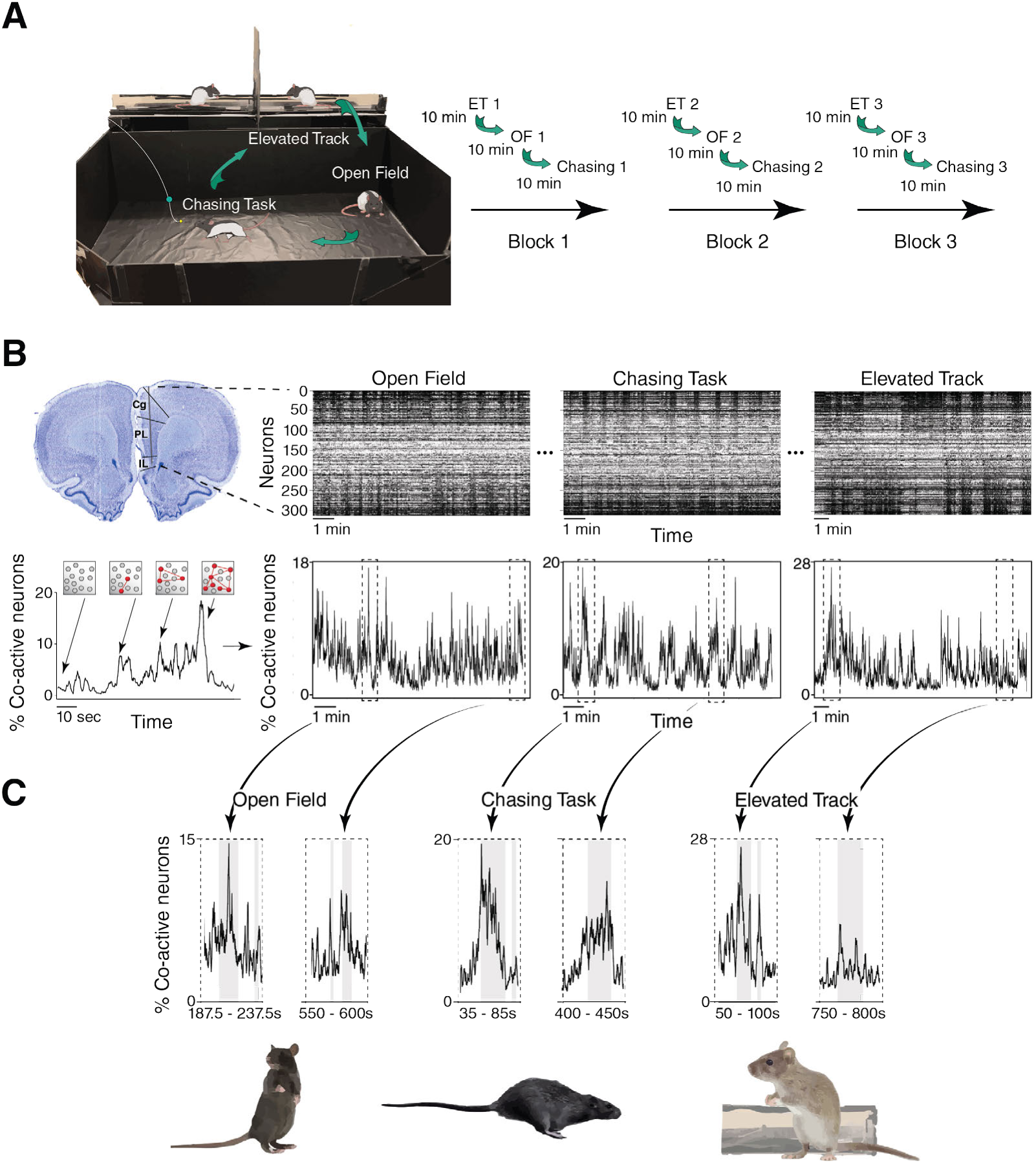
Naturalistic behaviors modulate prefrontal neural population structure. (A) Experimental design and timeline. Animals completed the same sequence of three tasks over three consecutive blocks for a total of 9 sessions. Each session lasted 10 minutes, and animals were switched directly from one task to the next (right). (B) (upper left) Nisslstained tissue section from an experimental animal showing the probe trajectory through Cg, PL and IL cortical regions. (upper right) Example raster plots for the same 313 neurons from one animal in the open field, chasing task and elevated track; units are sorted vertically based on PC loadings in each task. (lower left) A schematic illustrating how time-varying coactivity plots (bottom) reflect the fraction of coactive neurons within the population (above). (lower right) Fluctuations in neural coactivity in the spike rasters above. (C) (upper row) Examples of epochs with high coactivity in each task. (lower row) Illustrations of the behaviors the animals expressed during the highlighted peaks in coactivity.

The recording probes traversed mainly layer 5 of the cingulate (Cg), prelimbic (PL) and infralimbic (IL) regions (Paxinos & Watson, 2006) (Figure 1B, left; Figures S1-3), yielding a sum of 1711 well-isolated single units across tasks and animals. Since the behavioral tasks were not scripted, we used changes in the structure of population activity to signal salient behavioral events. We applied principle component analysis (PCA) and sorted spike rasters from each session by their PC loadings (Methods), which revealed abrupt shifts in spike train activity at different times in each task (Figure 1B middle, right). To better resolve how population activity changed during such events, we constructed a dynamic graph of spiking activity in each task by assigning an edge between pairs of neurons at each time point when they were strongly coactive (Figure 1B, lower left; Methods) (Bullmore & Sporns, 2009). By plotting the edge density (*i.e.* the fraction of coactive neurons) at each time point in each session (Figure 1B), we found epochs with high and low degrees of structure (Figure 1B, lower right), and used 3D tracking and video recordings to identify which actions occurred during those bouts. We note that we were limited to behaviors that could be identified based on the head and back, and that the analysis did not include the face, whiskers, limbs or tail.

Recurring subsets of behaviors such as rearing, chasing bait in the chasing task, or leaning on the wall of the linear track were associated with higher levels of neural coactivity, often marked by distinctive peaks, whereas sedentary behaviors such as grooming or eating in place were attended by drops in coactivity (Figure 1C). Data from one animal was used initially to label actions with ”high” coactivity (*i.e.*, when the fraction of coactive cells exceeded the 90th percentile of shuffled data for at least 5 of 21 bins; Methods) and ”low” coactivity (*i.e.*, coincident spiking *<* the 10th percentile of shuffled data for the same number of bins). This approach highlighted 7 distinct behaviors across the three tasks in the first animal, which were confirmed subsequently in the other two rats (Figures S4-9). In the open field these included “rearing”, “active investigation” and “head bobbing” (*i.e.* stationary eating or grooming). The same behaviors were expressed in the elevated track, which additionally had bouts of wall leaning (”front paws on wall”), “wall sitting” and “wall hanging”, and the chasing task included manually-scored chasing bouts. We did not find anatomical or topographical organization for the behavior state representations in any animal (Figures S1-3, right). To segment the behaviors systematically, we generated a supervised classifier based on tables of kinematic features that distinguished each behavior (summarized in Figures S4-9). For each animal and each task, the classifier was trained on a subset of tracking data and applied to the rest of the recordings, then user-adjusted with video recordings to validate each behavioral epoch. We also point out that the list of 7 behaviors was not exhaustive of the animals’ behavioral repertoire, but reflected actions which impacted neural coactivity consistently and could be identified readily.

### Single-cell and population coding of distinct behavior states

With each of the 7 behaviors time stamped in the timelines of each task (Figure 2A), we asked whether behavior-related coactivity fluctuations reflected global changes in firing rates or selective changes in subsets of cells. To answer this we binned the spike trains for each neuron in 50 ms bins and generated coactivity matrices for all possible pairs of neurons during each behavior (Methods). We then plotted the mean co-activation between each pair of neurons during each behavior, which produced 7 coactivity matrices that were unique to each behavior (Figure 2A, middle row). The activity patterns comprising each matrix emerged dynamically from background activity when the animals engaged in each behavior, such as chasing or rearing, and persisted until that behavior stopped and another began (Video S2).

**Figure 2.**
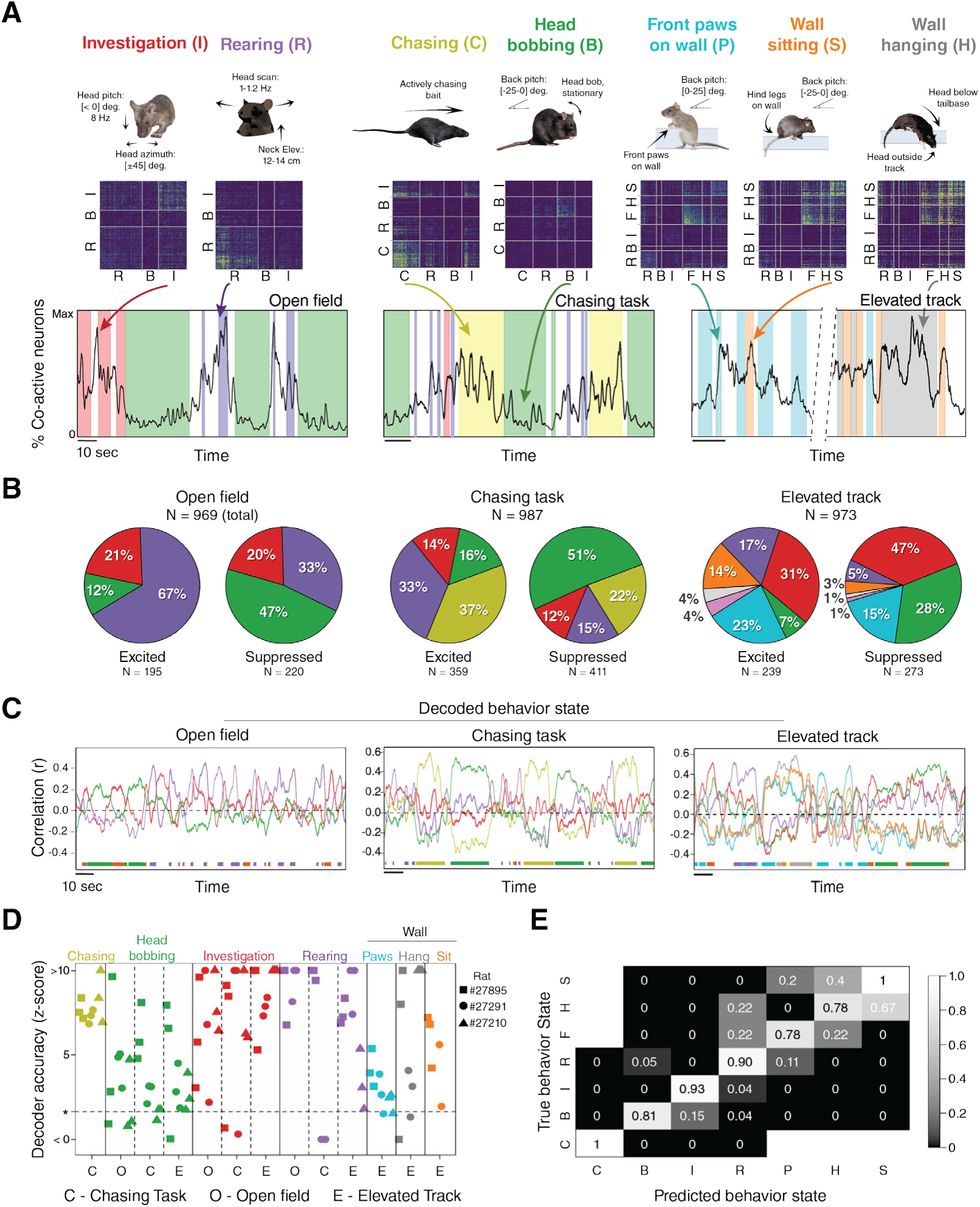
Neural ensemble representation of 7 behavior states. (A) (top row) Each of the seven behavior states, with core physical features indicated. (middle row) Mean pairwise coactivity matrices for each behavior state from an example animal (rat #27895, N = 313 cells) Highest to lowest coactivity is scaled from yellow to blue. (bottom row) Examples of coactivity fluctuations (color coded) during each of the behaviors, with neurons clustered based on their behavioral modulation in each task. (B) Proportions of excited and suppressed single units for each task, pooled across animals. 42.9% of 969 cells stably encoded behaviors in the open field (20.2% were stably excited, 22.7% suppressed); 79% of 987 showed tuning in the chasing task (36.6% excited, 42.4% suppressed); 52.7% of 973 units encoded behaviors in the elevated track (24.6% excited, 28.1% suppressed). Color code same as (A); magenta wedges in Elevated Track pie charts indicate socially- modulated cells. (C) Population vector decoding of behavior states across different sessions of like tasks. Colored lines indicate correlation values between population vector activity in the test session and the template for each behavior. Colored dots (bottom) indicate the expression of each behavior. (D) (left) Scatter plot showing z-scored decoding accuracy for each behavior from each animal in each task. Columns correspond to session types, colors indicate specific behaviors, symbols denote animals. Dashed line indicates a significance level of *α* = 0.05. (E) Proportion matrix summarizing the fraction of times a behavior was predicted beyond *α* = 0.05 in (D) plus false-positives.

Individual neurons were classified as behaviorally selective if their average firing rates stably increased or decreased during a given behavior in at least two sessions (Methods), with between 40 and 80% of single units passing this criterion in each task (Fig 2B; Methods). Different proportions of neurons represented different behavior states in each task, possibly reflecting their relevance in the different conditions. For example, ”rearing” and ”chasing” were the most common behaviors among excited neurons in the open field and chasing tasks, respectively, and head bobbing was the most common among suppressed neurons in both tasks, likely reflecting food consumption during foraging or after chasing (Figure 2B). The elevated track, on the other hand, presented the richest local environment of the three tasks, and ”active investigation” was the most commonly encoded behavior for both excited and suppressed neurons (Figure 2B). We also measured social proximity and orofacial interactions between animals on the elevated track, but these did not consistently modulate spiking activity so were not analyzed further. Within each task, behaviorally-excited neurons were exclusive to one behavior, and units excited during one behavior were typically suppressed during others (see also Jung et al., 1998; Mulder et al., 2003; Hyman et al., 2005). The suppression of neural firing when switching from preferred to non-preferred behaviors may have also contributed to rapid up-down dynamics in population coactivity through time (Figures 1B, 1C and 2A).

We next considered behavioral representation at the population level. We assessed this by training decoders on population vector activity during labeled behavior states in two sessions, and used those as templates to predict the occurrence of behavior states in the third session. Templates were generated using correlations between binned spike train activity during each behavior state in the first two sessions, and a running correlation was calculated between the template and the population vector activity in the test session (Methods). The running correlation predicted correct behaviors nearly continually throughout each test session (Figure 2C, line graphs; behaviors indicated by colored dots at bottom). The population codes for some behaviors appeared mutually exclusive (*e.g.*, a high probability of “chasing” meant a low probability for “head bobbing”, Figure 2C, middle), whereas others were more similar, especially for wall-directed behaviors in the elevated track (Figure 2C, right). Decoding accuracy was highest for chasing, rearing, and active investigation (Figure 2D), and lowest for head bobbing, likely due to the associated suppression of firing rates. Wall-directed behaviors on the elevated track, which elicited similar patterns of neural activity (Figure 2A) also had lower, but still above-chance decoding accuracy (Figures 2D and E).

Having tracked the kinematics of the animals’ heads in each task, we also identified allocentric head direction cells in each animal, confined primarily to prelimbic cortex (Figures S10 and S11; Methods). In sessions with sufficiently many simultaneously recorded cells, neural coverage of all directions was evident in the population activity (Figure S10B). Furthermore, the signals appeared to generalize across tasks, since a decoder trained on head direction signals in the open field could predict ongoing head direction in the other two tasks (Figure S10C), and directional tuning was maintained during different behavior states (Figure S10D), irrespective of the task in which the behavior occurred.

### Behavior states are not explained by low-level motor features

Since each behavior state comprised a collection of kinematic features, and prefrontal neurons often respond to multiple variables (Parthasarathy et al., 2017; Grunfeld et al., 2018; Aoi et al., 2020) including actions (Voloh et al., 2023), we asked whether the behavioral representations could be accounted for by one or more posture or movement primitives. We focused on chasing, rearing and active investigation since these behaviors had obvious kinematic components and were encoded robustly. We assessed the extent of overlap between cells that encoded a specific behavior state (*e.g.* chasing) as well as two core kinematic features used to classify that behavior (*e.g.* running speed and head azimuth velocity; Figure 3A; Figures S4, S6). Critically, we only included neurons that stably encoded their preferred features in both sessions in the analysis. From the chasing sessions, a sum of 386 units were modulated by chasing, running speed, head azimuthal velocity or combinations of these (Figure 3A, B, left). However, the majority of chasing-modulated neurons (142 of 251 chasing units; 37% of 386 total units; Figure 3B, left) encoded chasing only, and not running speed or head movement. In fact, substantially smaller fractions of cells were co-modulated by chasing and head velocity (42 cells, 11% of 386 total), chasing and running speed (79 cells, 20%), and only 12 cells (3%) were tuned to all three (Figure 3B, left). A hypergeometric test confirmed that the degree of overlap across the different categories was lower than expected by chance (p*<* 0.001; Methods). This was also the case for rearing and its constituent attributes of head pitch and neck elevation (Figure 3B, middle; p = 0.026, hypergeometic test), and for active investigation and its main components of head pitch and head azimuth (Figure 3B, right; p *<* 0.0001; data for individual animals shown in Figures S12-14).

**Figure 3.**
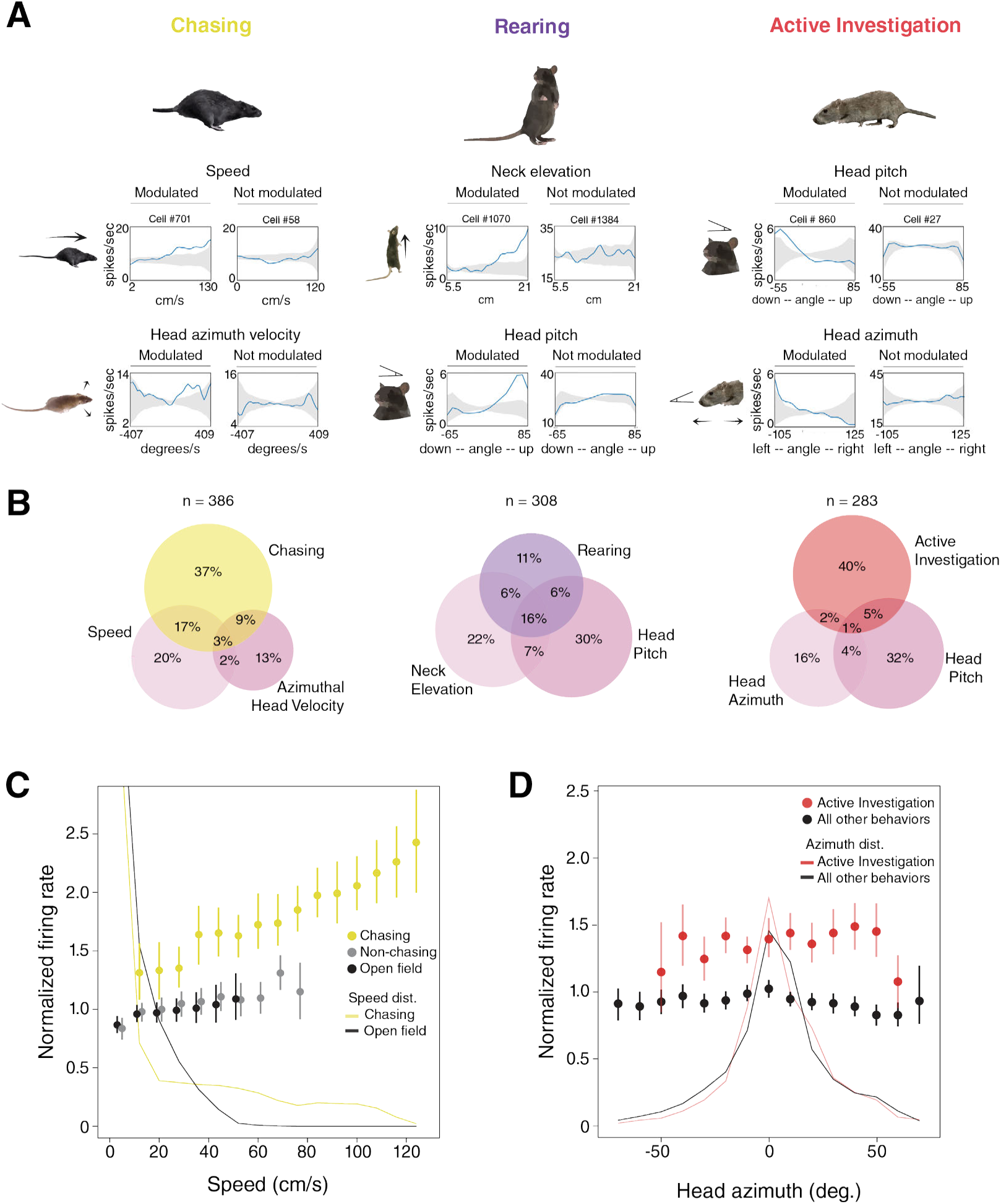
Behavior states are not accounted for by movement kinematics or running speed. (A) (left) Examples of two chasing-excited cells (#701 to the left and #58 to the right), with the former showing stable tuning to running speed and head movement, and the latter not. (Middle) Examples of two cells excited during “rearing”, with one (#1070, left) encoding neck elevation and head pitch, and the other (cell #1384, right) being unaffected. (Right) Examples of ”active investigation” cells that were modulated by the downward and leftward posture of the head (cell #860), or not (cell #27). (B) (left) Venn diagrams showing overlap of single units tuned to chasing, running speed or head azimuthal velocity in the chasing task; (middle) same, for single units stably tuned to rearing, neck elevation or head pitch (open field data used due to better sampling); (right) same, for cells stably tuned to active investigation, head azimuth or head pitch (chasing sessions used due to better sampling). Cell counts include both behavior-excited and -suppressed neurons. (C) The sensitivity of chasing-excited neurons to running speed was potentiated during chasing bouts compared to non-chasing intervals or during open field foraging (n = 51 chasing-excited neurons also active in open field). (D) Prefrontal neurons more strongly encoded forward head azimuth posture during active investigation than other behaviors (n = 54 behaviorally-excited cells active in all three tasks). For both (C) and (D), dots indicate the mean, error bars ±95% confidence interval (CI).

This was surprising *prima facie* since it was not clear how a cell could encode chasing behavior, for example, yet not be consistently modulated by running speed. However, closer inspection showed that neural responsiveness to kinematic variables depended on the state of behavior when they occurred. That is, the sensitivity of chasing-tuned neurons to speed was stronger while the animals chased the bait, with the same cells firing at lower rates for the same running speeds between chasing bouts or in the foraging task (Figure 3C). The same phenomenology was exemplified by ”active investigation” neurons, which more strongly encoded head azimuth angles between ±45 degrees during bouts of active investigation than other behaviors (Figure 3D). Neural ensembles also tended to form before and remain active after physical acts were made. Figure S15 illustrates this, where rearing-specific ensembles formed prior to rearing, when all paws were still on the floor, then transitioned smoothly to the chasing ensemble when the bait was introduced in the arena, but while the animal was still immobile. The neural ensembles for rearing could also remain active between bouts of rearing (Figure S16) whereas, in the chasing task, ”chasing” ensembles dominated even when animals were physically reared up to grab the bait at the end of a trial (Figure S17).

### Population activity evolves at slower timescales in mPFC than motor cortex

The propensity of mPFC neurons to anticipate and outlast the physical expression of their preferred behaviors prompted us to ask whether overall population dynamics in mPFC were inherently slower than other regions more directly linked to movement production, such as motor cortex. We tested this by generating autocorrelations of population coactivity from mPFC recordings in the open field and compared them against M1 recordings in foraging rats recorded previously within-lab (Mimica et al., 2023). The recordings were scaled temporally to match the kinematics and running speed of the animals across M1 and mPFC data sets (Figure S18; Methods), yet fluctuations in population coactivity were slower and more varied in mPFC than in M1 (Figure 4A). We quantified these differences by fitting a power law decay model for each recording, which confirmed in all cases that autocorrelation values in M1 dropped faster than in mPFC (decay parameter values, signifying a faster decay, ranged from 1.60-2.57 ±1.50-2.79 (lowest and highest values ±95% CI) in M1 versus 0.45-1.14 ±0.35-1.24 in mPFC; Methods). The slower fluctuations in mPFC are consistent with the interpretation that prefrontal neural ensembles evolve slower than in cortical areas that generate movement, possibly reflecting motivational states that drive actions rather than the actions themselves.

**Figure 4.**
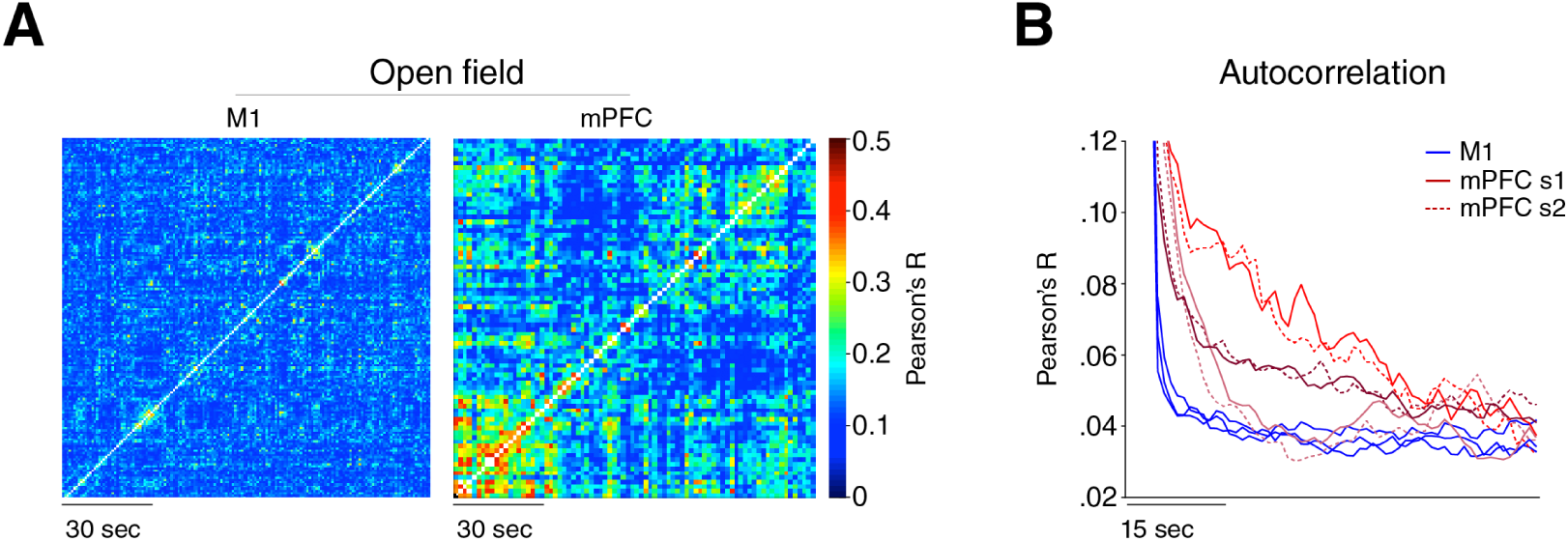
Population coactivity fluctuates more slowly in mPFC than in M1. (A) Autocorrelation matrices showing slower fluctuations in population coactivity in mPFC than primary motor cortex during open field foraging. (B) Autocorrelations of neural coactivity fluctuations in mPFC from each animal in two open field sessions (solid and stippled red lines, respectively) and single open field sessions in M1 (n = 3 rats, blue).

### Effects of task on behavioral selectivity

Up to this point we established that behavioral coding in mPFC was not anchored to motor execution, yet was stable across repeats of the same task, so we sought to determine the extent to which behavioral tuning was shaped by the tasks themselves. To do this we first assessed whether individual cells encoded the same behaviors expressed in each of the different tasks (rearing, head bobbing and active investigation), and only considered neurons which had stable tuning correlates across both sessions in the analysis. However, action coding in most cells did not carry over between unlike tasks (Figures 5A and S19; data from each animal shown in Figures S20-22), even though behaviors were defined identically (Figures S4-6) and the range of postural sampling was highly overlapping (Figure 5B; though dwell times for rearing were longer in the elevated track). Environmental geometry appeared relevant, since exploratory behaviors like rearing and active investigation were encoded by larger total common pools of cells in the open field and chasing tasks (Figure 5A; Figure S19), which were in the same arena, and less so across those tasks and the elevated track (Figure 5A; Figure S19). Although smaller fractions of cells encoded the same behaviors across tasks, we asked if this was nevertheless sufficient to reliably represent the same behaviors across them. We tested this using decoders trained on population vector activity from two tasks, then testing if they could predict the occurrence of the correct behavior in the third. We found indeed that actions could be decoded correctly irrespective of the task, and with only slightly lower accuracy than when decoding within-task, as illustrated by confusion matrices from a single rat (#27895; Figure 5C), and with data summed across all animals and tasks (Figure 5D). Prefrontal ensembles thus collectively retained behavioral fidelity despite considerable influences of task, which aligns with earlier findings that prefrontal ensembles stably encode confidence (Masset et al. 2020) and directed attention (Wu et al., 2017) in different tasks or environments.

**Figure 5.**
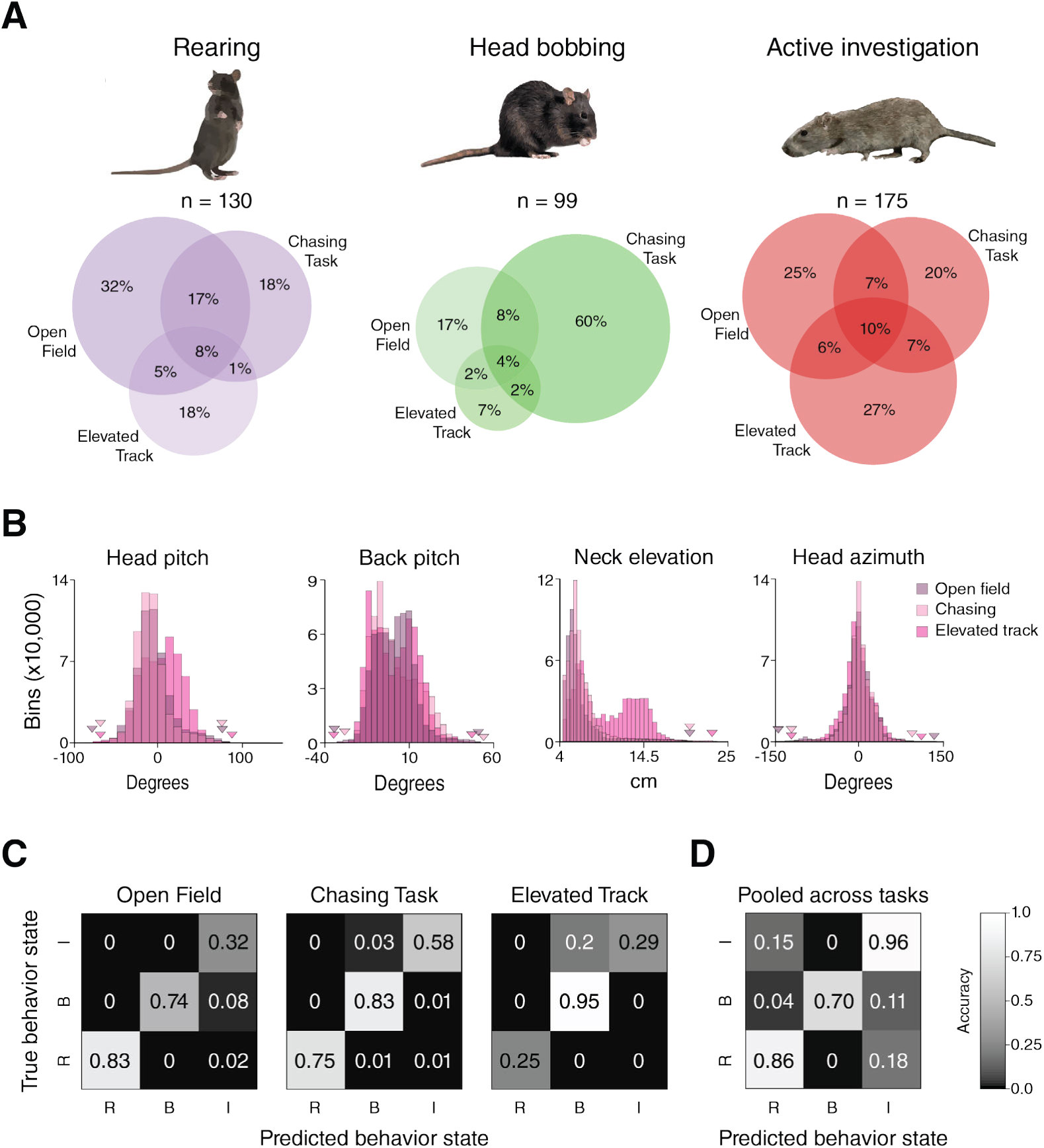
Behavioral selectivity varies across tasks at the single cell level but is maintained in neural ensembles. (A) (left) Venn diagram showing the overlap of neurons excited during rearing in each task, pooled across animals. (middle) Same, for head bobbing, and (right) the same for active investigation. (B) Histograms showing the ranges of kinematic sampling in each task (upper and lower bounds denoted by triangles). (C) (left) Confusion matrix showing the accuracy of a decoder, trained on population recordings from the chasing task and elevated track, at predicting corresponding behavior states in the open field. (middle) Same for the chasing task, where the decoder was trained on ensemble data from the open field and elevated track. (right) Same for the elevated track, with the decoder trained on ensemble data from the other two tasks. (D) Proportion matrix summarizing the fraction of instances when each behavior from each animal was decoded beyond a significance level of 0.05 across tasks (individual data points shown in Figure S23).

## Discussion

In the present study, we examined neural population dynamics in mPFC in freely moving rats performing minimally structured tasks, and found that mPFC neurons encoded a range of holistic, species-typical behaviors associated with information seeking, spatial exploration and mapping, pursuing prey, consuming food or grooming (Renner & Seltzer, 1991; Thinus-Blanc, 1996; Belzung, 1999; Lever et al., 2006; Butler et al., 2017; Layfield et al., 2023; Shan et al., 2023). We identified these behaviors using tasks that avoided stereotyped alignment of cognitive processes and actions, then performing an unbiased analysis of the structure of population activity. Unlike sensory and motor cortices (Parker et al., 2020; Schneider et al., 2020; Zimmermann et al., 2020; Mimica et al. 2023), action representation in prefrontal areas was not well accounted for by pose or movement, with the coding of kinematic features being apparently gated by the momentary behavior the animal expressed. The disconnection between prefrontal population activity and motor output was underscored by the fact that prefrontal population activity fluctuated at inherently slower timescales than M1. Because of this, we may have missed the behavioral coding phenomenology in mPFC had we used a more typical approach of segmenting behaviors from kinematics first then querying for neural correlates afterward (*e.g.* Markowitz et al., 2019; Mimica et al., 2023). Although the behavioral tuning of single cells was impacted by the task the animal performed, coding was stable at the ensemble level, indicating a capacity for flexible yet stable coding of behaviors in different circumstances. Our approach of tracking many features while making few *a priori* assumptions also led to the finding of prefrontal head direction cells, to our knowledge, for the first time reported in rodents. Together, our results suggest that prefrontal ensembles encode naturalistic behavior flexibly, in an anatomically distributed manner (Le Merre, et al., 2021), and at a level of abstraction that depends more on the task at hand than motor output. This supports the view that prefrontal areas flexibly define whatever neural state space is required for a given task or environment (Wilson et al., 2014; Sharpe et al., 2019).

Because the behavioral tasks were unstructured, we relied on changes in population coactivity to signal events that engaged prefrontal regions, without pre-supposing which features or which level of behavioral organization was important (Anderson & Perona, 2014; Gris et al., 2017; Krakauer et al., 2017; Berman, 2018; Buzsáki, 2020). This reduced experimenter bias, and the behavior states it allowed us to identify, such as active investigation and rearing, could reflect the occurrence of cognitive processes related to planning or decision making as implemented in naturalistic settings. In this case, our findings would complement a considerable history of work showing the formation of neural ensembles during trial-based cognitive tasks (Miller, 2000; Ridderinkof et al., 2004; Gilbert & Burgess, 2008; Fuster, 2015; Hanganu-Opatz et al., 2023), but here couched in freely-expressed actions spanning sub-second to seconds-long scales. By trading off the predictability of a learned task structure, it also becomes possible to establish how prefrontal cortices steer action selection in a manner that avoids biases caused by prior training (Arlt et al., 2022; Peters et al., 2022) or restricting experiments to a single context (Lee et al., 2022).

A growing number of studies have also used naturalistic designs to probe mPFC tuning in relation to unrestrained behavior. One line of work focused on the game of hide and seek in rats (Reinhold et al., 2019) where, similar to our study, the authors (Bagi et al., 2022) first characterized the structure of the neural data before linking it back to behavior (Little & Rubin, 2019; Buzsáki, 2020). Two key differences between that study and ours were that hide and seek has an inherent structure which shapes behavior and requires training, and that behavioral classification in that work used video-based tracking that could not distinguish task variables from body kinematics. Tracking kinematic variables proved critical in earlier work investigating prefrontal coding of navigational sequences, for example, where what initially appeared as neural correlates of working memory were better accounted for by stereotypic head movements during certain trajectories (Euston & McNaughton, 2006). Even more recent work investigated action representation directly in prefrontal regions of freely behaving macaques, and found that neural firing along adjacent prefrontal regions encoded naturalistic actions and the transitions between them (Voloh et. al., 2023). Because of the deep intertwining of action and cognition (Fine & Hayden, 2021; Kane et al., 2023), it was by virtue of recording the same cells in multiple tasks in our study that we could show that low-level kinematic variables were not hard-coded, but were elements of complex medleys of features evoked when animals engaged in motivated behaviors.

Another aspect of behavioral representation we identified was tuning to allocentric head direction, which was stable across behavioral tasks. This adds to a growing literature indicating a role for prefrontal cortices in spatial navigation (Euston et al., 2012; Patai & Spiers, 2021; Maisson et al., 2022), which is intuitive considering the ubiquity of navigation in many goal-directed natural behaviors. New work in freely moving macaques, for example, demonstrated that prefrontal neurons represent spatial features such as place and head direction alongside task-related behavioral variables, like choice and reward, as well as self-paced actions (Maisson et al., 2023). Orbitofrontal cortex in rats was also shown to be critical for mapping and remembering the spatial location of food rewards (Feierstein et al., 2006; Basu et al., 2021), and exhibiting spatial firing fields modulated by extrinsic features such as the flavor of available food (Wikenheiser et al., 2021). Based on the anatomy and connectivity in rodents it is most likely head direction signals are not computed locally in prefrontal areas but are inherited from other regions. These could include hippocampal (Ferino et al., 1987), retrosplenial (Vogt & Miller, 1983; Van Eden et al., 1992) or thalamic inputs (Ferino et al., 1987; Jankowski et al., 2014; Ito et al., 2015), though future work is needed to define their respective contributions, or if there is an anatomical organization underlying where prefrontal head-direction cells are expressed or not (see Poucet, 1997).

In summary, we found that prefrontal population activity in freely moving rats is shaped by momentary, holistic behavioral states rather than single features or functions. The composite nature of the representations implies that they emerge from a confluence of inputs from different cerebral structures, which is consistent with existing frameworks for prefrontal cognitive function in general (Fuster, 2001; Miller & Cohen, 2001; Hanganu-Opatz et al., 2023). Discerning the contribution of different inputs to the selection and expression of specific behavior states will require further measurements and manipulations of select neural pathways, as done with dopaminergic signaling in relation to motivated behaviors in song birds (Roeser et al., 2023) and spontaneous action in mice (Markowitz et al., 2023). The longer timescales over which behavior state representations were expressed, and their mercurial linkage with kinematics, make a purely motoric explanation unlikely, particularly in light of inherent differences in population dynamics with M1. Because of this, it was by studying the structure of neural coactivity first, rather than movement kinematics, that we were able to identify clear behavioral representation in mPFC. By considering the dynamic density of connections between nodes in large networks (Bullmore & Sporns, 2009), graph theoretic methods as used here could be applied to expose behavioral representation in any number of systems in the brain, during wakefulness or sleep, or as a consequence of learning.

## Author contributions

Conceptualization: B.A.D, I.V.R., F.N., J.R.W.; Methodology: I.V.R., F.N., B.A.D., J.R.W.; Formal analysis and investigation: I.V.R., F.N., I.H.; Project administration: I.V.R., B.A.D, J.R.W.; Funding acquisition: B.A.D, J.R.W.; Resources: B.A.D, J.R.W.; Supervision: B.A.D, J.R.W.; Writing - original draft preparation: I.V.R.; Writing - review and editing: I.V.R., B.A.D, F.N., J.R.W.

## Acknowledgements

We thank M. Andresen, K. Haugen, A.M. Amundsgård, P. J. B. Girão, and H. Waade for technical and IT assistance; S. Eggen for veterinary oversight; M. P. Witter, G.M Olsen, David Klindt and current and alumni members of the Whitlock and Moser lab for helpful discussions.

## Funding

This work was supported by a Research Council of Norway FRIPRO grant (No. 300709) to J.R.W., a Research Council of Norway grant (iMOD, No. 325114) to B.A.D., an NTNU Medical Faculty Fellowship (RSO) to I.V.R., the Centre of Excellence scheme of the Research Council of Norway (Centre for Algorithms in the Cortex (CAC), grant No. 332640), the National Infrastructure scheme of the Research Council of Norway – NORBRAIN (grant No.197467), and The Kavli Foundation.

## Compliance with Ethical Standards

The authors declare that they have no conflict of interest.

## Data availability

The datasets generated during the current study are available here: https://figshare.com/s/f09f8ac2cf9fff1f8f35. The codes used in the present study are available here: https://github.com/fredrine/behavior_state_mPFC_scripts

## Methods

### Animals

All experiments were performed in accordance with the Norwegian Animal Welfare Act, the European Convention for the Protection of Vertebrate Animals used for Experimental and Other Scientific Purposes, and the local veterinary authority at the Norwegian University of Science and Technology. All experiments were approved by the Norwegian Food Safety Authority (Mattilsynet; protocol ID # 25094). Three female rats (age: 11-15 weeks, weight: 285-335 grams at time of surgery) were used in this study. Two additional animals were also recorded from, but their data was discarded due to poor tracking and/or quality of acquired neural data.

All rats were housed in familial groups in enriched cages prior to surgery and whenever possible housed with their experimental partners after surgery. If any risk to implant damage was detected, the implanted animals would be housed individually in plexiglass cages (45 x 44 x 30 cm). Animals had *ad libitum* access to food and water throughout the entire study and were housed in a temperature and humidity-controlled environment on a reversed 12h light/12h dark cycle. Animals were handled daily from at least 8 weeks old.

No animals were food or water deprived before, during or after the experiments.

### Surgery

Animals were anesthetized initially with 5% Isofluorane vapor mixed with oxygen and mainted at 1-3% throughout the surgical procedure. When anesthetized, they were placed in a stereotaxic frame and injected subcutaneously with Metacam (2.5mg/kg) and Temgesic (0.05mg/kg) as general analgesics. Body temperature was maintained at 37°C with a heating pad. A local anesthetic (Marcain 0.5%) was injected under the scalp before the first surgical incision was made, followed by clearing the skull of skin and fascia. Following the clearing of tissue, adjustments were made to make sure the skull was leveled before a high-speed dental drill was used to open a craniotomy over the medial prefrontal cortex and to drill multiple holes in the skull for inserting jeweler’s screws, used later to anchor dental acrylic (Kulzer GmbH, Germany). A single jeweler’s screw was inserted over the cerebellum as a ground and reference screw for the probe implant, connected with a silver wire and sealed with SCP03B (silver conductive paint). Animals were subcutaneously administered fluids at the start, at the half-way point of surgery and immediately post-surgery.

A single-shanked silicon probe (Neuropixels version 1.0, IMEC, Belgium) was stereotaxically lowered in the right hemisphere of all three animals at coordinates AP: +2.5-3, ML: 0.5-0.6, DV: 4-4.5. The craniotomy and unimplanted length of the probe were enclosed with a silicone elastomer (Kwik-Sil, World Precision Instruments Inc., USA). After allowing the silicone elastomer to harden, a thin layer of Super-Bond (Sun Medical Co., Ltd., Japan) was applied to it and the skull to increase bonding strength between the skull surface and dental acrylic. To minimize light-induced electrical interference during the experiments the first layer of dental cement was dyed black and care was taken to completely cover any light-sensitive parts of the probe. Undyed cement was further applied to secure the probe. A plastic screw-top casing made from a 15-ml Falcon centrifuge tube was used to encase the implant outside the brain, serving both as a protective housing and as a base for attaching the rigid body for tracking the head in 3D during recordings. The tube was also secured to the skull with undyed cement. Once the cement had cured, sharp edges were removed with the dental drill at the end of surgery. Following surgery, the animals were placed in a heated (32°C) chamber to recover. Once awake and ambulating, they were returned to their home cage and allowed food and rest before data collection commenced.

### Recording procedures and behavior

Electrophysiological recordings were performed using Neuropixels 1.0 acquisition hardware: A National Instruments PXIe-1071 chassis and PXI-6133 I/O module for recording analog and digital inputs (Jun et al. 2017). Implanted probes were operationally connected via a headstage circuit board and interface cable above the head. An elastic string was used to counterbalance excess cable, allowing animals to move freely through the entire recording area during recordings. Data acquisition was performed using SpikeGLX software (SpikeGLX, Janelia Research Campus), with the amplifier gain for AP channels set to 500x, LFP channels set to 250x, an external reference and AP filter cut at 300 Hz. Spike activity was recorded and collected from bank 0, *i.e.*, the most distal 384 recording sites.

Behavioral experiments and neural recordings where performed a minimum of 2.5 hours after animals had fully recovered from surgery.

Animals were habituated for at least 3 sessions in all three tasks before surgery, but care was taken not to over-habituate the animals, keeping them as naive as possible to the tasks and surroundings, but without causing them stress or anxiety. It was most important to ascertain the animals’ pre-surgical inclination to the Chasing Task, as most animals had an inherent drive to chase after the bait, but a minority of animals did not. This trait was crucial to identify in the animals prior to assigning them to surgery, as there was no extrinsic motivation to chase since the animals were not food deprived at any point. If the animal did not show a proclivity for instinctual chasing, the animal was excluded from further consideration as an experimental subject. Additionally, one ”dry-run” of the experimental task-sequence, including transitions between tasks in the appropriate order, was performed before surgery to ensure they had at least one experience with transitions between the tasks and contexts.

### 3D Tracking and labeling

Animals had circular marker cut outs of 3M retro-reflective tape (Opti-Track, catalog no. MSC 1040; Natural Point Inc., Corvallis, OR, USA) placed on their body at three standard points (neck, back and tailbase) that were shaved during surgery to allow for tracking of the trunk in 3D. To capture head position and movement a rigid body with four retro-reflective spheres, sizes 7.9 mm, 9.5 mm and 12.7 mm (OptiTrack, catalog no. MKR079M3-10, MKR095M3-10 and MKR127M3-10, respectively), was constructed. During the recording experiments, the rigid body was attached to the centrifuge tube cemented in the implant to allow for tracking of the head while keeping the probe protected. The 3D position of each marker was recorded using 11 near-infrared (NIR) recording cameras (10 NIR, and one black/white camera was used for referencing; Flex 13 cameras, Part number: FL13, OptiTrack, Oregon, USA) which tracked the XYZ position of each marker at 120 FPS in optical motion capture software (Motive version 2.2.0; OptiTrack). 8 cameras were angled down from the ceiling, while the remaining 3 cameras were angled more horizontally at a lower height to capture behavior on the elevated track. For synchronizing the tracking and neural data streams, three LEDs were affixed to the sides of the recording arena and were also captured by the motion capture system. The LEDs were controlled by an Arduino Microcontroller with C++ code to generate unique sequences of irregularly timed digital pulses (250ms duration of each flash; random 250 ms ≤ IPI ≤ 1.5 sec) that were transmitted to the different acquisition systems throughout the recordings.

The individual markers were labeled using built-in functions in Motive on raw .tak files. After recordings were completed, 3D data was reconstructed by creating a marker set for the three body markers, a rigid body construction for the capture of the animal’s head (in the sessions with two animals, two separate and unique rigid body constructs were used to identify the animals) and a rigid body was constructed to capture the synchronizing LEDs. Using Motive’s built-in ”autolabel” function, markers were identified and labeled, with the remaining unlabeled markers hand labeled by the experimenter. Errors in marker assignments were also corrected during this manual process.

For frames where markers were occluded (by the animal’s body or physical objects) or not detected by the tracking cameras (but markers were still available on reference video), reconstruction of tracked point was attempted using in-built functions in the Motive software. Different modes of reconstruction were used depending on the duration of occlusion, the behavior of the animal over the frames of relevance, the identity of the marker itself and the potential for validating the reconstructed markers. Specifically, methods of interpolation used for reconstruction were ”cubic”, ”constant”, ”linear”, and ”pattern based”.

After reconstruction and labelling of the markers, the positional information from each session was exported as a .csv-file with units expressed in meters and individual markers as outputs. A custom-written Python script was used to further process and align tracking information with the synchronization stream into a .pkl file. This file was then loaded in a custom graphical user interface (GUI) to construct a head coordinate system and merge the tracking data with spike times into a final .mat file used for subsequent analyses. More details on the procedure are described in (Mimica et al. 2018). The minimum threshold for determining the animals’ running speed was 5 cm/s.

### Behavioral tasks

All three tasks were performed by the animal in a structured sequence, back-to-back, which included a minimum of three repeats for all tasks. The task-sequence was always (1) Elevated Track, (2) Open Field, (3) Chasing Task. After (3), the animal was placed back on the elevated track, and a second animal (serving as a social cue) was placed on the other end of the track and the experimental task-sequence would restart. The animals were moved by the experimenter from the elevated track to the arena and back during brief pauses in data acquisition to avoid introducing electrical noise by having physical contact. Each session lasted 10 minutes, with one session comprising one ”behavioral task”.

#### Elevated Track

The elevated track was designed to provide a behavioral context containing a social element with which the recorded animal was free to interact through a physical separating barrier. The track was elevated 64 cm high above the arena floor. The track itself was 10 cm wide with 11 cm high walls, and was 190 cm long, with a barrier in the middle (width: 61 cm, height: 53 cm) separating the two animals. The barrier had an opening (width: 10-17cm, height: 22cm) cut into it where it met the track to allow the animals to interact, but not climb on each other. The opening itself was covered with a plastic grid and strips of fine mesh to keep the rigid body on either animal from getting stuck.

Both animals were tracked in 3D, but electrophysiological recordings were made only from the experimental animal during the elevated track sessions. Data acquisition and tracking started once both animals were placed on either side of the barrier. Animals were allowed to move freely on their side of the track and interact in the middle through the barrier of their own volition. At times the experimenter sought to encourage the animals to approach each other by throwing food crumbs close to the barrier, but the animals chose whether to approach or not.

#### Open Field

After the elevated track, animals were placed in an open arena (125cm x 190cm x 50cm) beneath and beside the elevated track, and allowed to explore it and forage for food. There were no salient cues placed on the arena walls, but as there were no curtains outside the arena, the animals could use extra-arena cues in the recording room for orientation. The animals were allowed to roam freely until the session time had elapsed and the chasing task commenced. We used this condition in effect as a baseline since there were no task-relevant variables to encourage specific behaviors.

#### Chasing Task

In the chasing task, the experimenter attached a small piece of food to the end of a fishing line, which hung from a fishing rod that the experimenter weaved in semi-random patterns to encourage the animal to chase the bait, but in a winding fashion, across the entire arena space. A retroreflective spherical marker was attached 9 cm above the bait so it could be tracked during chasing trials. The time point from when the animal pushed off with its hind legs in the direction of the bait, to the time point of when the animal caught the food was defined as a single trial. Once the animal had the food, the experimenter waited until the animal had either finished eating or lost interest and moved away from the food. Some animals routinely ate the food where they caught it, while others took the food to a corner in the arena to consume it. After the animal had either finished eating or lost interest in the catch, the experimenter waited a few seconds before initiating a new chasing trial. This procedure was repeated over the course of the 10 min session. When the session time elapsed the animal was either transferred back to the elevated track to begin a new task sequence, or the experiment concluded if the animal showed signs of fatigue, stress or non-compliance during the latest task-sequence.

### Spike sorting, cluster identification and unit profiles

All recording sessions for each animal were concatenated into a single binary file, one for each animal, to retain the identity of the recorded clusters across all tasks. Spike sorting was done using Kilosort 3 (Pachitariu et al., 2023), and all spike data was curated manually using Phy 2.0 to separate noise from single and multi-unit activity. Using custom-made code with metrics as described in Hill et al (2011) and adapted from https://github.com/AllenInstitute/ecephys_spike_sorting, single units were defined as well isolated if they had an ISI violation of ≤ 2 % in a *<* 1.5ms ±interval around the occurrence of a putative spike. Putative multi-units were separated from noise based on their waveforms. After manual curation, well-isolated single units were allocated to their respective sessions/tasks, and only units with ≥ 100 spikes per session were kept for further analyses. Putative duplicate spikes were removed and all final units were visually inspected as a final quality check using Phy. Furthermore, good units were split into fast-spiking (FS) and regular-spiking (RS) subtypes by performing K-means clustering (where k=2) on spike width, peak-to-trough ratio, full width at half maximum and hyperpolarization (or end) slope data (Niell & Stryker, 2008).

### Unit classification

#### Behavior classification

Using manually assigned frames that contained the behaviors of interest in a subset of sessions, we built a classifier using a multinomial regression, fitted using LASSO regularization, to create tables that described each of the manually labeled behaviors using information extracted from the 3D-tracking of the animals. These tables were then used to predict the likelihood that a given behavior was executed in each frame of the recordings, based on the tracking information of the animal. Behaviors that lasted less than 60 consecutive frames (≈ 0.5 seconds) where not included for further analysis. Behaviors that were too similar for the classifier to accurately separate (due to limitations in the detail and features included in the tracking) were assigned further restrictions or requirements manually. For example, it was not possible to have the requirement that the front paws were physically touching the top of the walls in the elevated track since the paws not tracked. The final set of frames used in analyses were inspected visually and corrected by the experimenter before proceeding to the analysis stage.

There were two behaviors that were not defined using this approach: the chasing behavior and social proximity. As noted above, the time points for each chasing trial were defined by manually investigating the videos and defining start time from the frame where the animal pushed off with its hind legs in the direction of the bait, and the end point was the frame where the animal caught the bait with its paws. One such chasing bout was defined as a single trial, and needed to last ≥ 500 ms. ”Social proximity” was defined as the frames when the animals where within 40 cm of each other on the elevated track, using the position of both animals from the 3D tracking.

#### Head Direction Units

Using tracking data for the animal’s head, head direction was calculated in the coordinate system of the tracking software, which defined 2D head direction in space before anchoring it to a digital landmark in the room. Since the animals were tracked at 120 FPS, each recorded frame was ≈ 8.33 ms, and head direction in each frame was binned into 10°bins for a total of 36 bins. The average firing rate per bin was calculated as the total number of spikes per bin, divided by the total time spent in that particular bin. All smoothed 1D rate maps were constructed with a Gaussian filter with a standard deviation of 1 bin, and only bins with a minimum occupancy of 200 ms were used for subsequent analysis. Head direction rate maps which had tuning peaks that exceeded the 95th quantile of the shuffled distribution for three consecutive bins were investigated further for directionality tuning. The shuffled distribution was generated by shifting the neural activity 1000 times in intervals ranging in length from ±10 seconds to the entire length of the session. For those units with head direction maps exceeding the shuffled data as described above, we calculated their Reyleigh vector lengths in a minimum of two sessions. Those cells which had Reyleigh vector lengths longer than 0.2 were then tested for stability. If the Pearson’s correlation for tuning curves from two separate sessions had p-values that exceeded the 95th percentile of the shuffled distribution for that unit, it was classified as a head direction unit.

Isomap was used only for visualization, using stable head direction cells from animal # 27895 (Diana), since that animal had a sufficiently large population of simultaneously recorded head direction cells for the visualization. For this, the Isomap parameters used were sigma = 30, minimum 100 spikes in a given session, downsampled to every 25th bin, n-neighbors = 30, n-components = 3.

#### Tuning to other postural features

Postural features of the head were expressed in Euler angles, were binned in 5°increments, whereas postural features of the back were in 2.5°bins. Neck elevation was binned in 1 cm bins. For all rate maps, the average firing rate per bin was calculated as the total number of spikes per bin, divided by total time spent in the bin.

### Histology

At the conclusion of the experiment, animals were deeply anesthetized with Isofluorane and injected intraperitoneally with an overdose of pentobarbital (Exagon vet., 400 mg/ml, Richter Pharma Ag, Austria), after which they were intracardially perfused with saline or Ringers solution, followed by 4% paraformaldehyde. Animals were then decapitated, and skin and muscle were removed from the skull before leaving it to post-fix overnight in 4% paraformaldehyde. The probe shanks were left in the brains to allow for better detection of the probe track during histological sectioning.

The following day the brains were extracted from the skull and stored in dimethyl sulfoxide (DMSO) at 4°C. On the day of sectioning, the brains were removed from DMSO, frozen with dry ice and sectioned with a sliding microtome at 30 *µ* m in the coronal plane in three series. The first series was mounted immediately on glass slides, Nissl-stained and cover slipped, and the other two series were kept in DMSO at -25°C and studied if the probe track was inconclusive in the first series. Using a digital scanner and scanning software (Carl Zeiss AS, Oslo, Norway), all brain sections were digitized and the images were visualized with Zen (blue edition) software and subsequently used to confirm probe placement and depth reconstruction. Exact locations of recording sites were estimated based on surgical coordinates, probe geometry and configuration, and local anatomical landmarks (Paxinos & Watson, 2006).

### Data analysis and statistics

#### Assigning behavioral modulation for single cells

To determine whether the activity of a cell was significantly modulated by a specific behavior state or kinematic feature, we used a modified version of a *t*-test. The mean spike count in bins with a specific behavior label, scaled by the standard deviation, was compared to that of bins without the label. Whereas this difference could be compared to the *t*-distribution if each bin was independent, we constructed a shuffled distribution to account for the temporal structure of the data. This was done by resampling the intervals of the specified behavior uniformly across the session while avoiding overlap, and calculating the difference in mean spike counts. This procedure was repeated *N* = 1000 times, and the corresponding *p*-value is calculated as 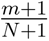, where *m* is the number of values in the shuffled distribution that exceeded the originally observed value. This was repeated for each type of behavior considered for the session type, then, for each cell, this set of *p*-values across behaviors was adjusted with a Benjamini/Hochberg correction to account for multiple behaviors being tested simultaneously. Ultimately the cell was considered up- or down-modulated during the behavior if the corresponding adjusted *p*-value was smaller than 0.05.

For population vector decoding, pair-wise coactivity analysis and UMAP projections (all described below) the n (n=313 for animal # 27895, n=215 for animal # 27291, n=75 for animal # 27210) cells that fired in all 9 sessions (elevated track 1-3, chasing 1-3, open field 1-3) were considered, and each spike train was binned at 50 ms.

#### Population vector decoding

For population vector decoding, the binned spike trains were smoothed using a Gaussian filter with standard deviation equal to 10 bins. For normalization each value in each spike train was replaced by its corresponding quantile from the distribution of values in that smoothed spike train (resulting in an even distribution of values between 0 and 1 for each cell).

For decoding within session type, two of the sessions were used to construct a decoder for the third. For decoding in open field session 1, for instance, the decoding procedure for each behavior *b* in {”rearing”, ”head bobbing”, ”active investigation”} was as follows:

1. For behavior *b* we calculated a vector ”thumbprint” *t_b_*, as the average population vector over time bins labeled as behavior *b* in open field sessions 2 and 3.
2. We calculated the correlation between *t_b_* and the population vector for each time bin in open field sessions 2 and 3.
3. We trained a logistic regression model on open field sessions 2 and 3 with this correlation as the covariate and a binary response equal to 1 for time bins labeled *b* and 0 otherwise.
4. We selected a cutoff maximizing Youden’s J statistic (= sensitivity + specificity - 1) (Youden, 1950), and calculated the corresponding cutoff, *c_b_*, in terms of correlation to *t_b_*.
5. We calculated the correlation between the population vectors in open field session 1 and *t_b_*.
6. If the correlation between the population vector at time bin *i* is larger than *c_b_*, the decoder classified that time bin as behavior *b*.

Note: if for a time bin *i* the thumbprint correlations for multiple behaviors were larger than their corresponding cutoff values, the decoder was set to classify the time bin by the behavior among these that had the largest thumbprint correlation. If no behaviors passed the threshold, no behavior was predicted. For decoding across session types, the procedure was similar, except that the decoder was made using all 6 sessions that were not of the same type.

#### Decoder accuracy

To evaluate the performance of the decoder and to determine whether the decoder accuracies were higher than what one would expect from chance, a chance level decoder was made. When making guesses for open field session 1, for instance, the chance level decoder was given the labels from open field sessions 2 and 3, similar to the actual decoder. It guessed randomly from these labels by using a circular shift of the combined vector of labels at a random lag and cutting the resulting vector off so that it had a length equal to open field session 1. This was repeated 10 000 times, and a confusion matrix was calculated each time. This yielded an empirical distribution for each entry in the confusion matrix, to which the observed matrix for the actual decoder was compared, by z-scoring these values using the mean and standard deviation of the empirical distribution. The z-scores are summarized in the scatter plots in Figures 2 and S23, and the fractions of how many times the z-scored value was beyond the 95th percentile of the standard normal distribution across 9 sessions and 3 animals are summarized in Figure 2D.

As for across-task decoding, the confusion matrices in Figure 5 show the proportion of time bins of a given behavior that the decoder correctly classified that behavior. The diagonal thus shows the proportion of time bins in that behavior that were correctly classified by the decoder. Note that the occurrence of each behavior is unbalanced, as some behaviors happen more often than others. Thus there is a bias in the decoder accuracies. Due to low sampling of some of these behaviors, downsampling the other behaviors to balance the data was not feasible. Instead, we constructed a method to evaluate the decoder which accounts for the imbalance.

#### Pairwise coactivity analysis

For the pairwise coactivity analysis, the binned spike trains were smoothed using a Gaussian filter with standard deviation equal to 5 bins. For normalization, each value in each spike train was replaced by its corresponding quantile from the distribution of values in that smoothed spike train. A binary activity vector for each cell describing whether or not the cell was active at a coarser time scale was then constructed by letting its value at time bin *i* be 1 if at least 5 bins in the 21 bin window {*i* − 10*, …, i, …, i* + 10} of the quantiled version was larger than 0.90.

For two cells *x* and *y* let *A_x_* and *A_y_* be the sets of indices for which their respective activity vectors have the value 1, and *n* be the length of the session (in number of time bins). We define the scaled session-wide coactivity as:

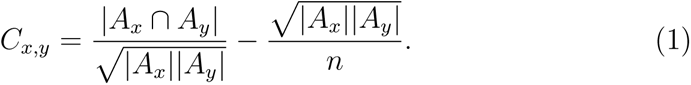

To determine which pairs of cells were significantly coactive, we constructed a shuffled distribution of scaled coactivity scores. We circularly shifted the activity vectors independently for each cell, and recalculated the session-wide coactivity for each session, selecting a threshold for significance at the 95th percentile of the combined shuffled distribution for all nine sessions. For animal # 27895, for example, this yielded a threshold of 0.067, and out of 48 828 possible pairs of cells (17.1, 16.8, 19.7, 16.9, 19.1, 17.0, 21.6, 22.1, 20.9)% were significantly coactive in (O1, O2, O3, C1, C2, C3, E1, E2, E3), respectively.

For the dynamic coactivity matrices, the entry corresponding to the pair of cell *i* and cell *j* at time *t* is 1 if the pair is significantly coactive in the session, and both cells are active at time *t*, and was 0 otherwise. The graph showing the proportion of coactivity over time is at time *t* defined as the proportion of significantly coactive pairs, where both cells are active at time *t*.

The ordering of cells in the dynamic coactivity matrix was constructed by first ordering the cells according to the relevant ensembles for that session type (e.g. ”chasing”, ”rearing”, ”active investigation” and ”head bobbing” for the chasing sessions). Then each of the remaining cells not belonging to an ensemble were grouped with the ensemble for which the average session-wide coactivity was the highest.

#### UMAP projections

For the UMAP projections the binned spike trains were smoothed using a Gaussian filter with standard deviation equal to 5 bins, and standardized by subtracting the mean and dividing by the standard deviation of the values in the smoothed spike train. They were then downsampled to every tenth bin to obtain a less correlated set of values and to reduce computation time. Two concatenated sessions (two chasing sessions, two open field sessions or two elevated track sessions) were used to construct the UMAP projections.

For UMAP projections highlighting specific behaviors, the out-of-behavior time bins were further downsampled randomly to match the same number of bins for within-behavior segments. For other projections, all time bins remaining after the initial downsampling were used.

The UMAP function (in R) was used with parameters: metric = “cosine”, n neighbors = 100 and min dist = 0.01.

### Software tools

Analyses were done in R and in the Python programming language. The following libraries in Python were extensively used: *scipy*, *matplotlib*, *pandas*, *numpy*, *pingouin*. In R the following packages were used: *stats*, *ROCit*, *umap*.

## Supplementary Video Legends

**Video S1. Behavioral tasks used.** Black and white video showing segments of (i) the elevated track, (ii) the open field foraging task, and (iii) the chasing task. The elevated track was located above the north wall of the arena, and the foraging and chasing tasks were performed in the same arena.

**Video S2. Dynamic emergence of distinct prefrontal ensembles during four behaviors in the chasing task.** (Left) Animation of tracking data from ∼1 minute of a chasing session; the bait marker is shown as a white circle. (Right top) Coactivity matrix with black dots appearing when pairs of neurons were coactive beyond threshold. Neurons were ordered based on their behavioral modulation by chasing (C), rearing (R), head-bobbing (B) or active investigation (I). (Right middle) Timeline showing the correlation between actual population vector activity and population vector templates for chasing (yellow), head bobbing (green), active investigation (red) and rearing (blue). Color-coded squares were overlaid on the coactivity matrix (right top) depending on which template best matched the population activity. (Right bottom) Fraction of coactive active neurons among all neurons in the recording (n = 313).

---

**Figure S1.**
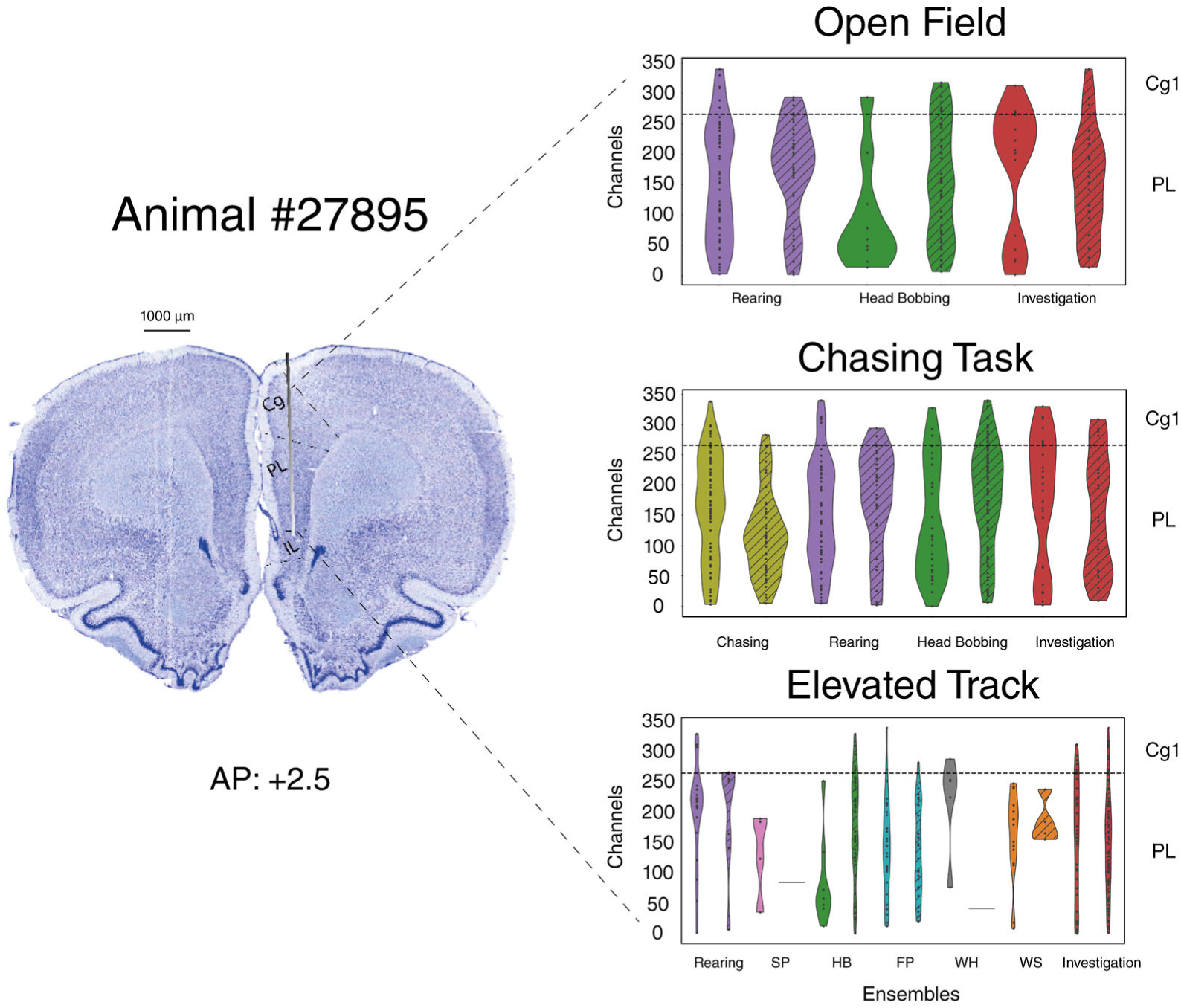
Histology showing implant location and distribution of behaviorally-tuned units along the probe for animal #27895. (Left) Nissl-stained tissue section showing the Neuropixel probe trajectory across the deep layers of the cingulate and prelimbic cortices. AP coordinate relative to Bregma is shown beneath. (Right) Color-coded violin plots depicting the recording channels at which behaviorally-excited (no fill) and -suppressed (stripe fill) cells were recorded in each task.

**Figure S2.**
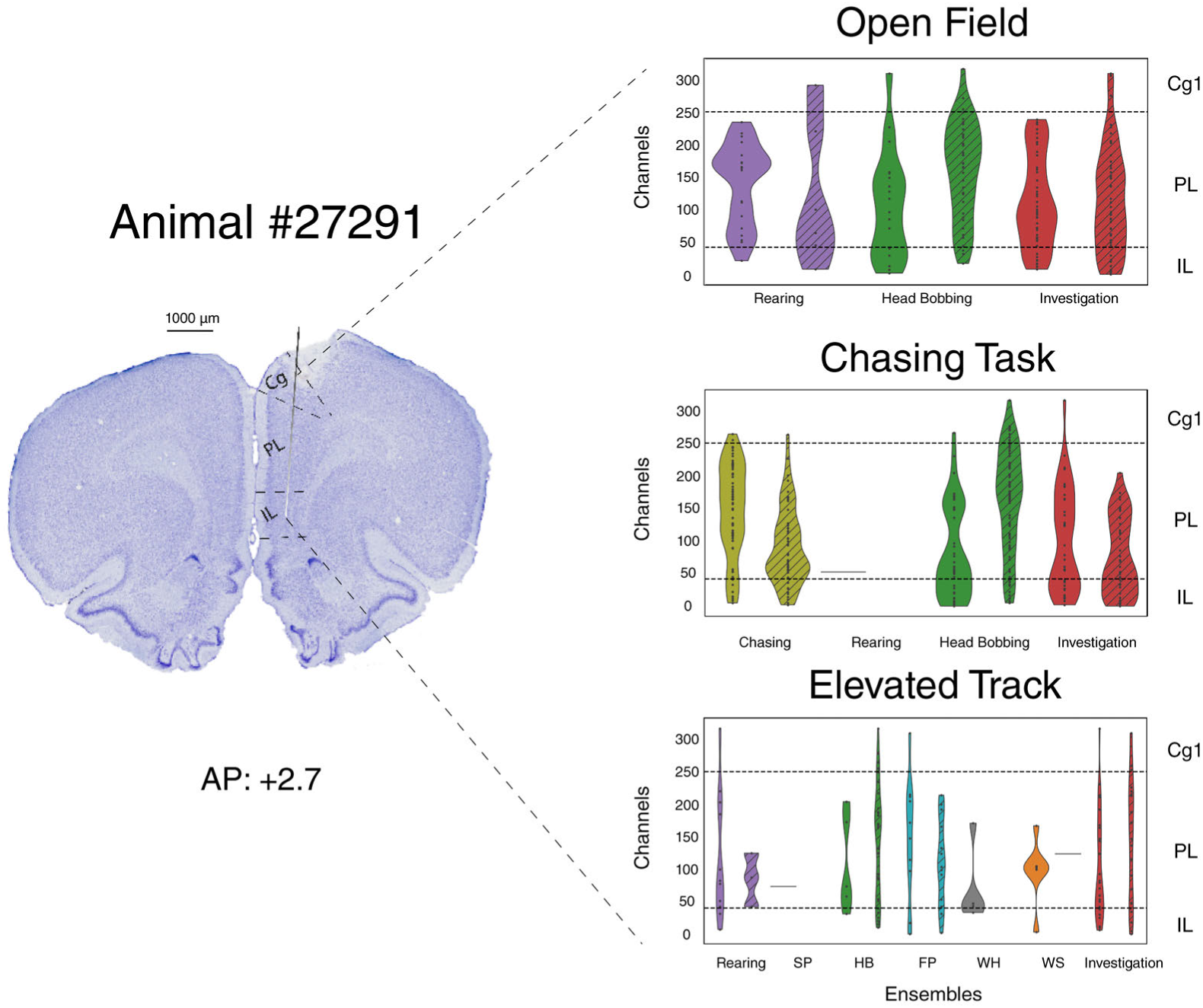
Histology showing implant location and distribution of tuned units along the probe for animal #27291. (Left) Nissl-stained tissue section showing the Neuropixel probe trajectory through the deep layers of the cingulate, prelimbic and infralimbic cortices. (Right) Violin plots show the same as for Figure S1.

**Figure S3.**
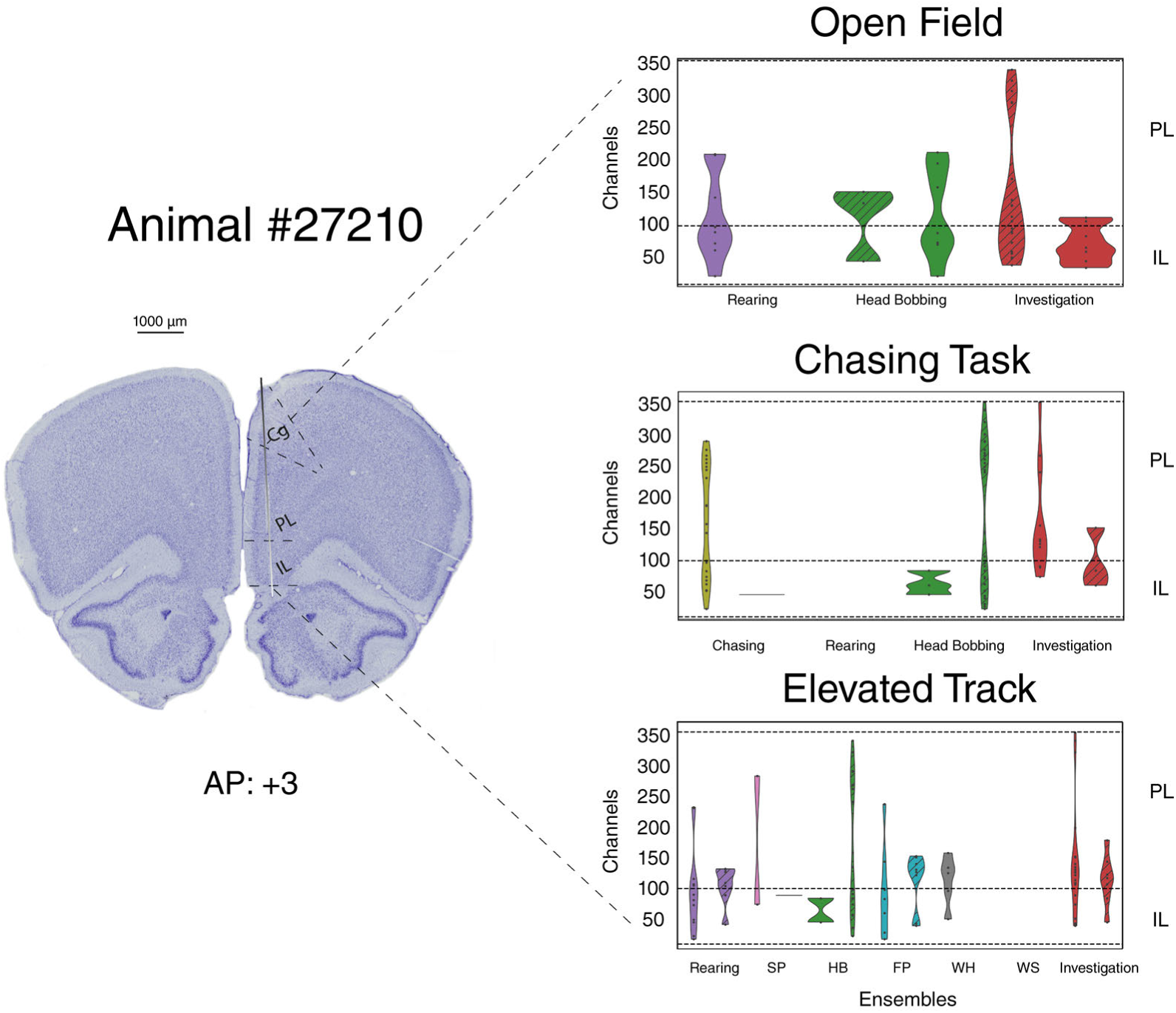
Histology showing implant location and distribution of tuned units along the probe for animal #27210. Same as Figure S2, with recording sites confined to deep layers in pre- and infralimbic cortices.

**Figure S4.**
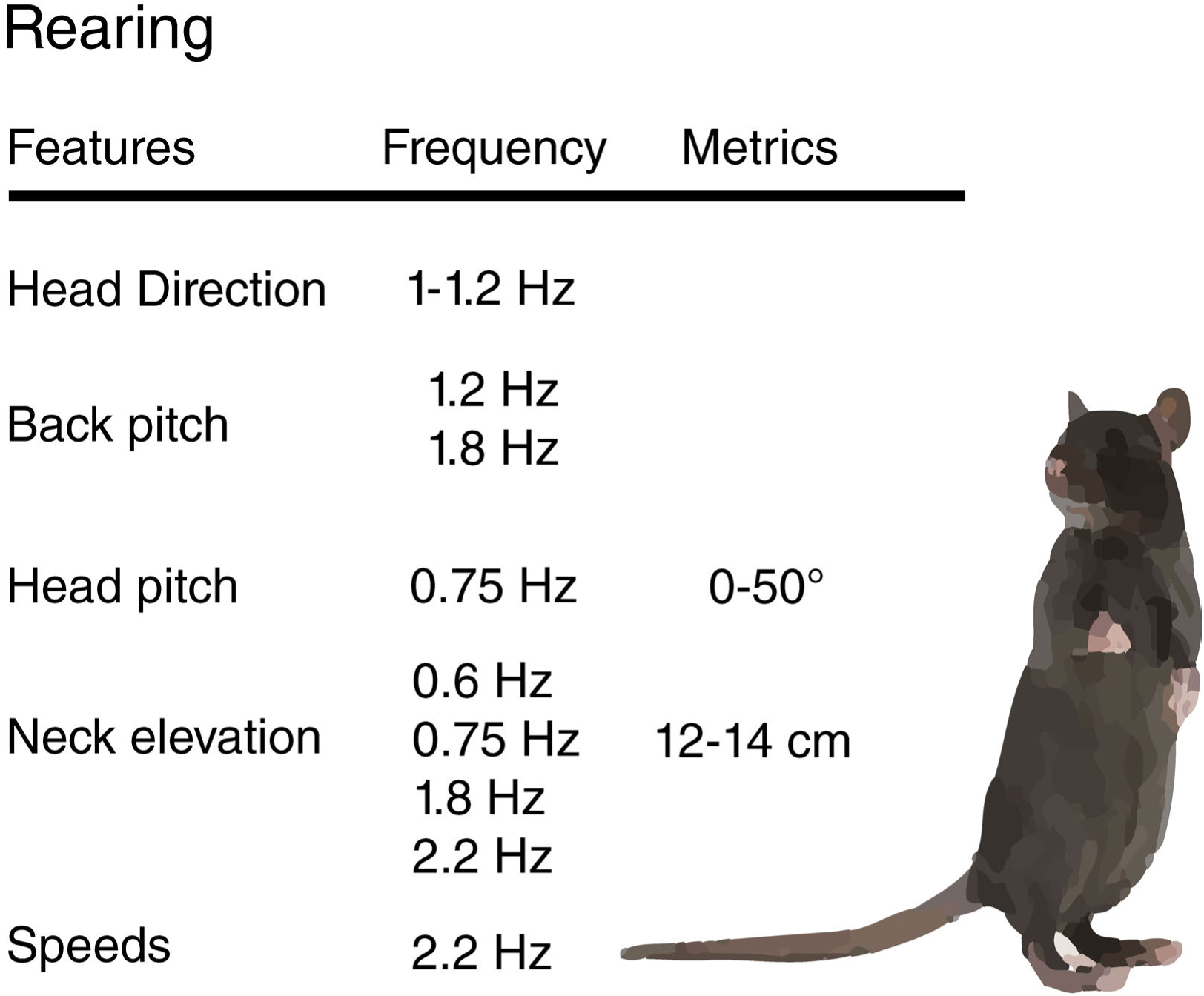
Summary of kinematic features defining ”Rearing” behavior as identified by the supervised classifier. For these and subsequent summaries: ”Features” refers to physical covariates tracked for the animals and the local recording area; ”Frequency” refers to the characteristic oscillation speeds of each feature, where relevant; ”Metrics” indicates angles or distances from the floor or between body markers.

**Figure S5.**
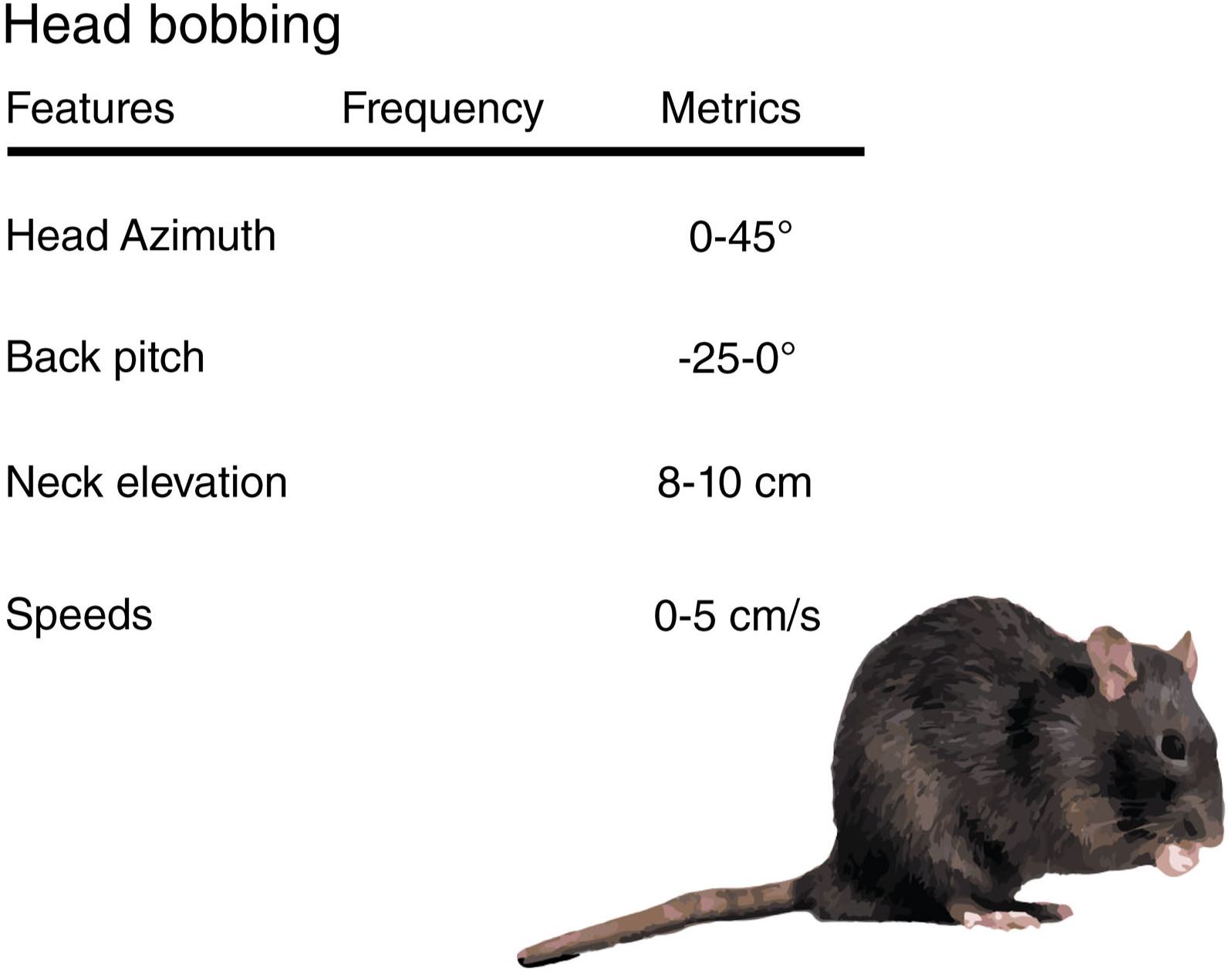
”Head bobbing” characteristics. Summary of kinematic features defining ”Head bobbing” as identified by the supervised classifier.

**Figure S6.**
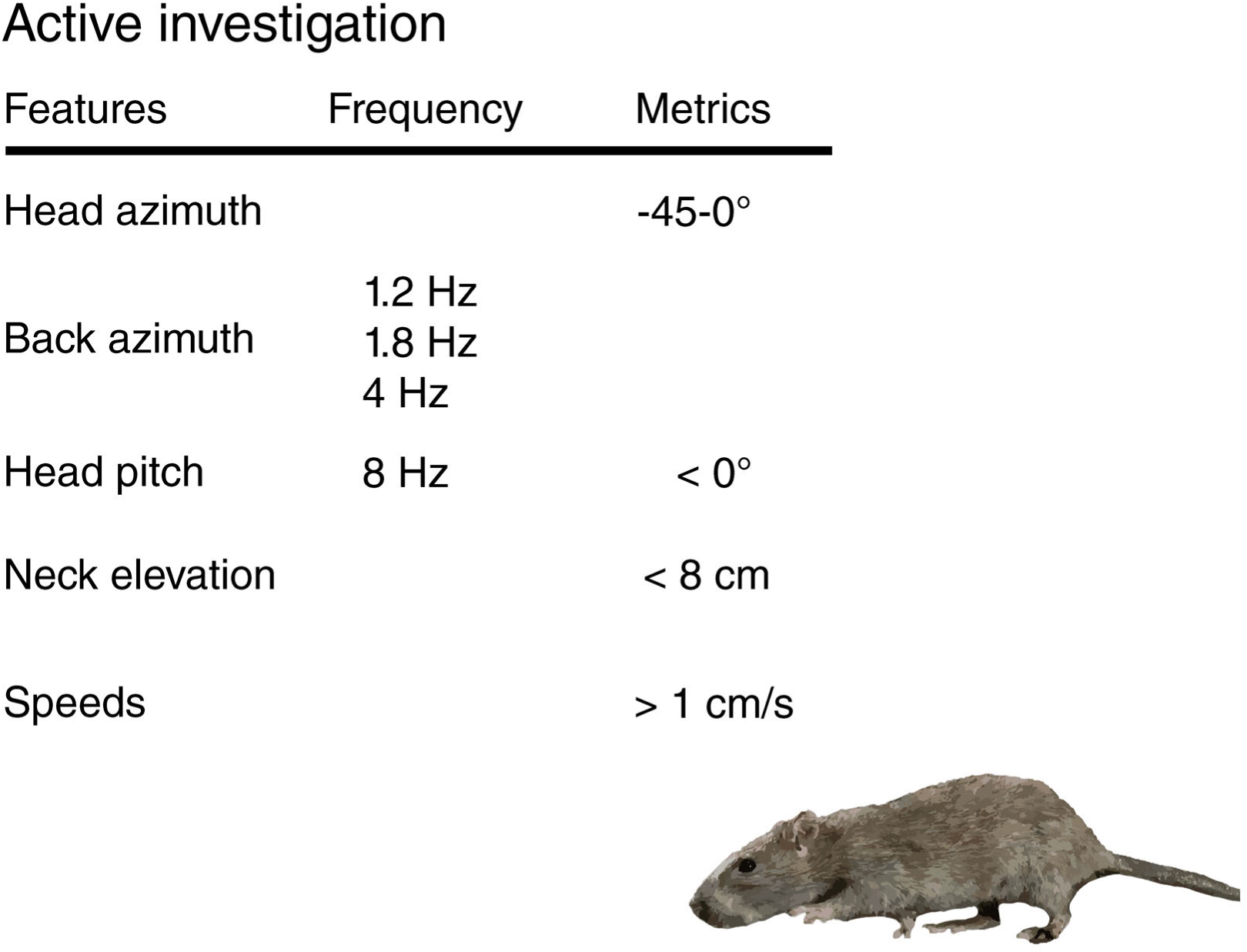
”Active investigation” characteristics. Summary of kinematic features defining ”Active investigation” as identified by the supervised classifier.

**Figure S7.**
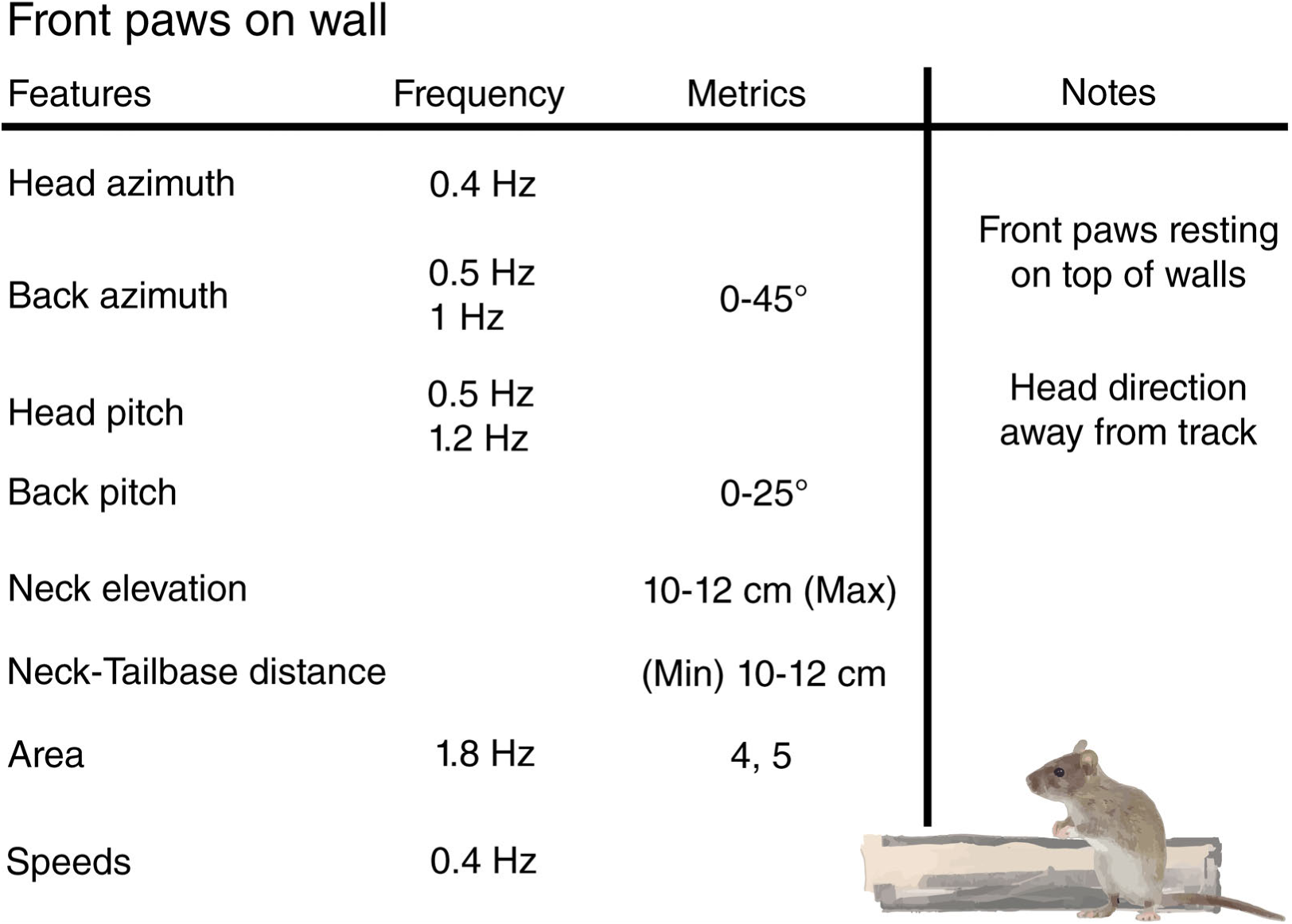
”Front paws on wall” characteristics. Summary of kinematic features defining ”Front paws on wall” as identified by the supervised classifier. ”Notes” refers to additional classification criteria used during video inspection; ”Area” indicates sections on the elevated track that could be physically occupied by the animal.

**Figure S8.**
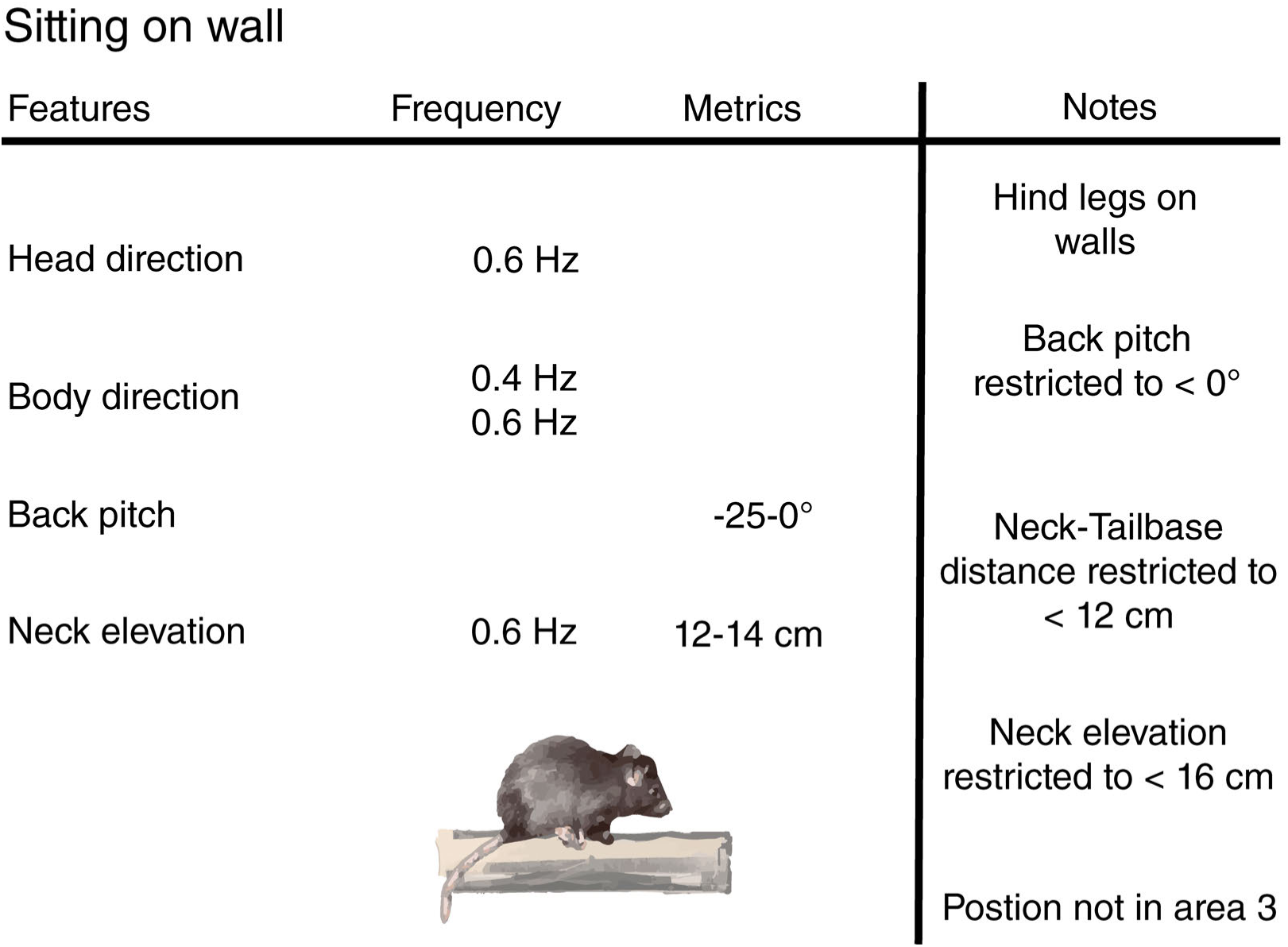
”Sitting on wall” characteristics. Summary of kinematic features defining ”Sitting on wall” as identified by the supervised classifier.

**Figure S9.**
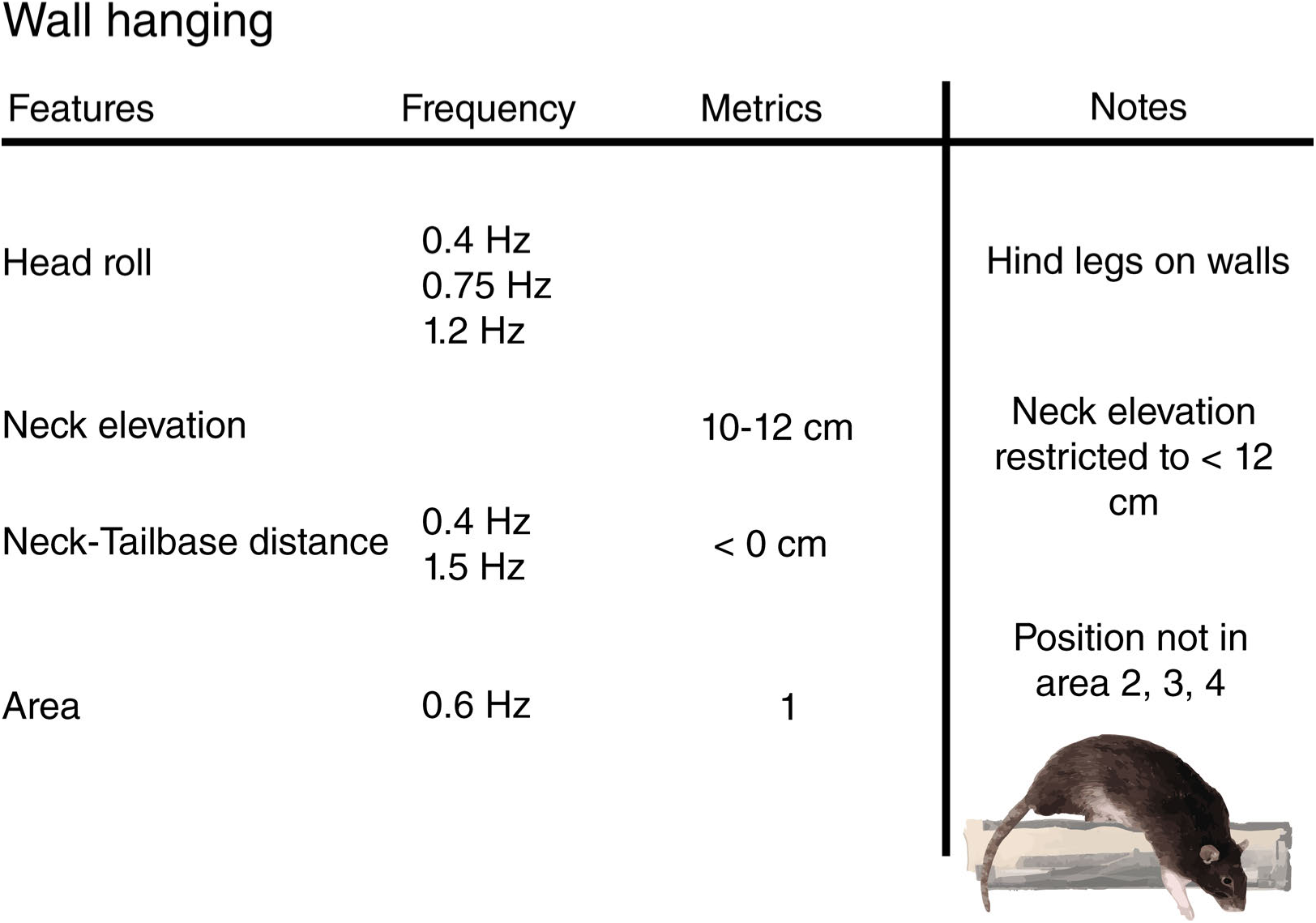
”Wall hanging” characteristics. Summary of kinematic features defining ”Wall hanging” as identified by the supervised classifier.

**Figure S10.**
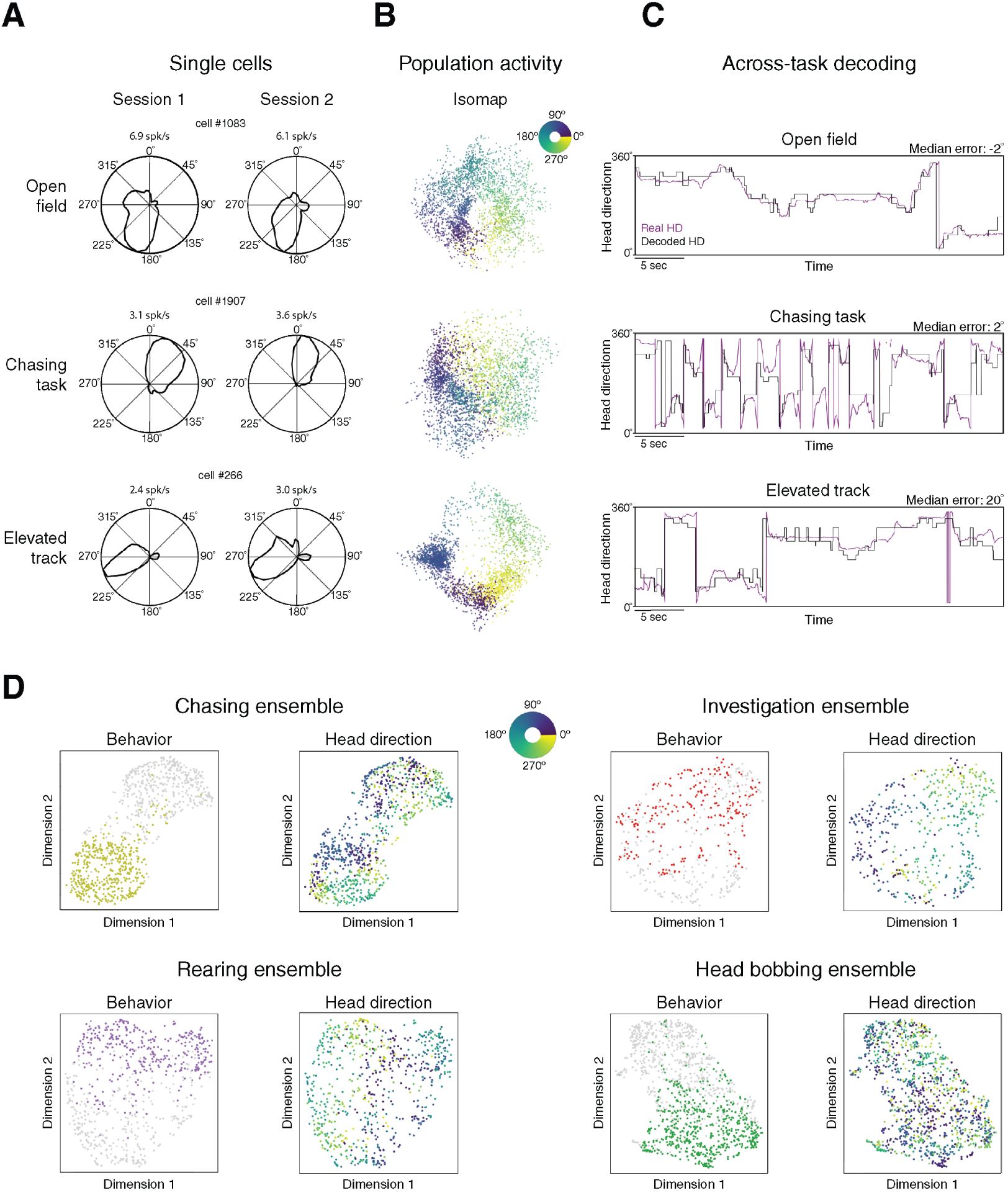
Head direction signals in prelimbic cortex generalize across tasks. (A) Examples of head direction cells in prelimbic cortex, with stable directional tuning across multiple sessions in each task. (B), Isomap visualization of population structure for stably tuned head direction units in each task. (C), Decoding of head direction in each task using a decoder trained on head direction signals from the open field. Purple lines show real head direction, black lines indicate decoded head direction. The smallest error was between the open field sessions, and the largest between the open field and elevated track. (D), Population activity of behaviorally tuned ensembles projected into UMAP-space. (upper left) UMAPs of chasing-tuned neurons. (left panel) Olive colored dots indicate time points when the animal was pursuing the bait in the chasing task; grey dots were intervals outside of chasing. (right panel) The same data colored according to the head direction of the animal at each time point, showing that the structure of directional tuning was maintained in the ensemble. (upper right) Same as upper left, but for “active investigation” ensembles in the open field task. Within-behavior time points are shown as red dots in the UMAP plot on the left, and the same data are color-coded for head direction to the right. (lower left) Same as the upper panels, but for “rearing” in the open field task; within-behavior time points are purple in the left UMAP; head direction is shown in the right. (lower right) Same, but for “head bobbing” ensembles in the chasing task colored in green for within-behavior time points (left), with (right) an apparent loss of directional structure in head direction signals in the ensemble.

**Figure S11.**
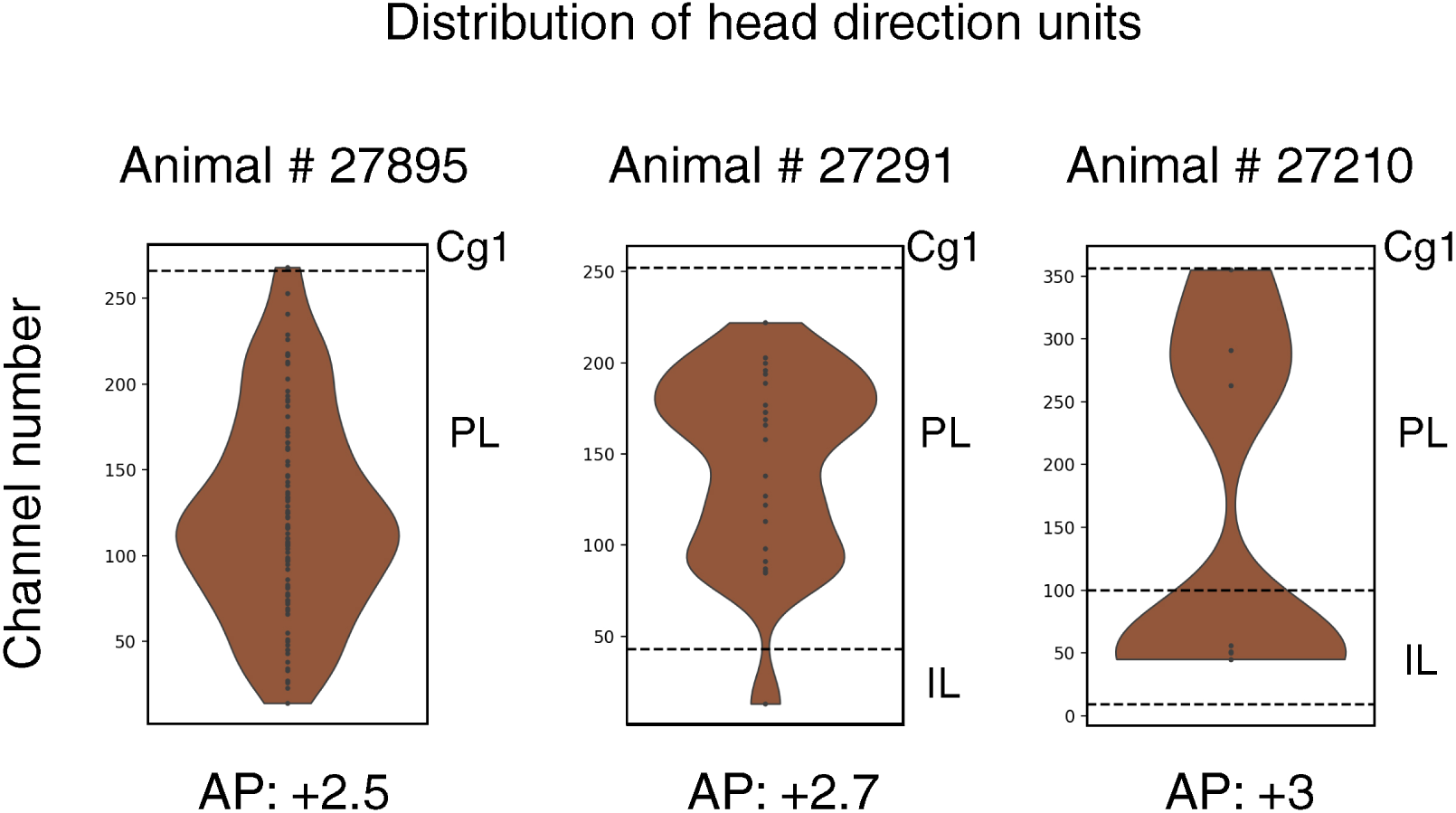
Distribution of head cells in prefrontal sub-areas in each animal. Violin plots showing the prevalence of head direction cells across the recording sites of each probe in each rat (95 cells in animal #27895, and 22 and 7 in #27291 and #27210, respectively). The vast majority of head direction cells were recorded in the deeper layers of prelimbic cortex, and at farther posterior implant locations. Anatomical boundaries marked with stippled lines.

**Figure S12.**
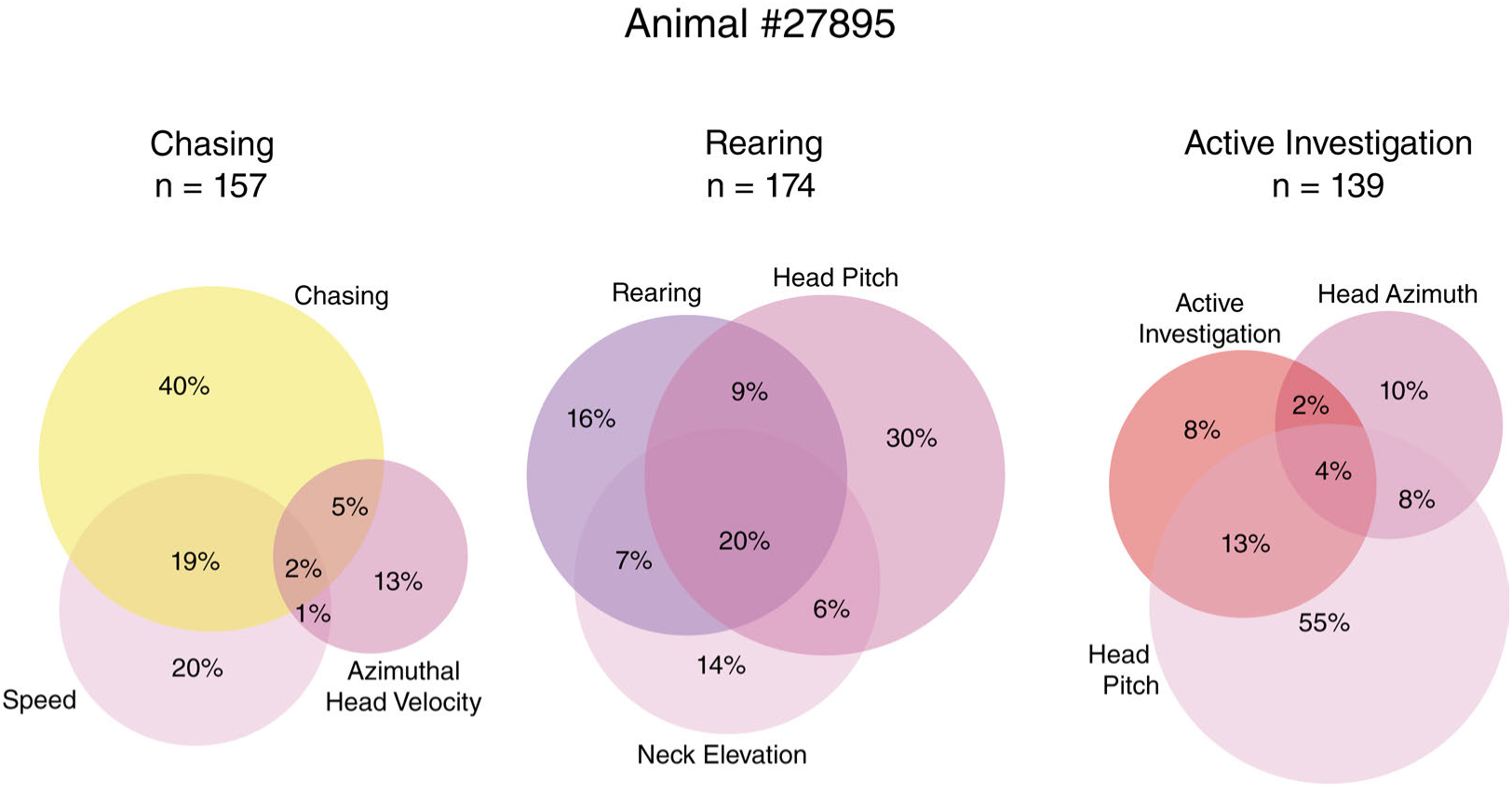
Overlap between units tuned to specific behavior states *versus* elementary kinematic features for animal #27895. (left) Venn diagrams showing overlap between single units that were stably tuned to the “chasing” behavior state, to running speed, or azimuthal head velocity (*i.e.*, the elementary physical features of chasing). (middle) Same, for single units stably tuned to the “rearing” behavior state, to neck elevation or head pitch. (right) Same, for cells stably tuned to “Active investigation”, to head azimuth or head pitch.

**Figure S13.**
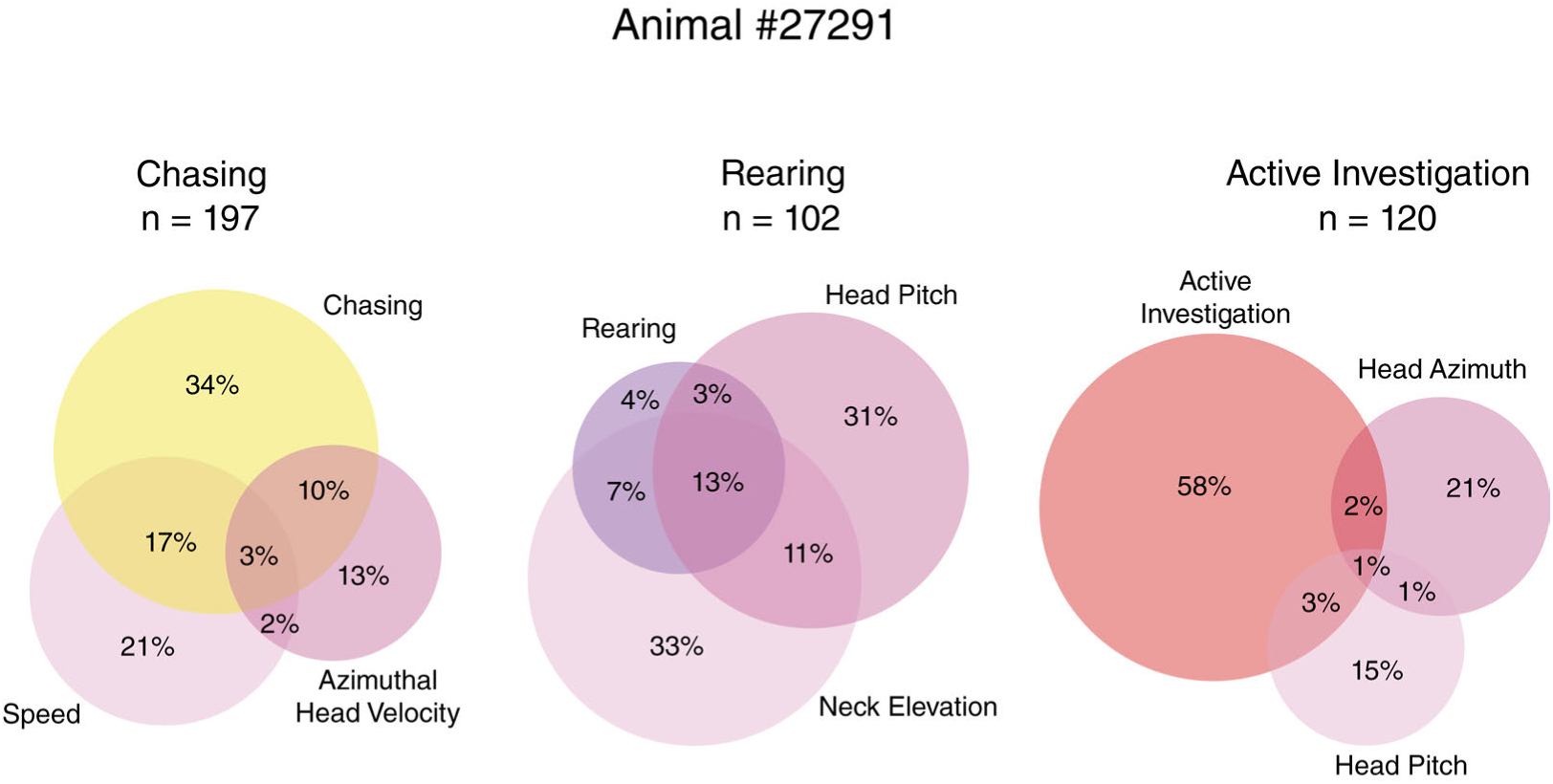
Overlap between units tuned to specific behavior states versus elementary kinematic features for animal #27291. Same as for Figure S12, but for the 2^nd^ animal in the study.

**Figure S14.**
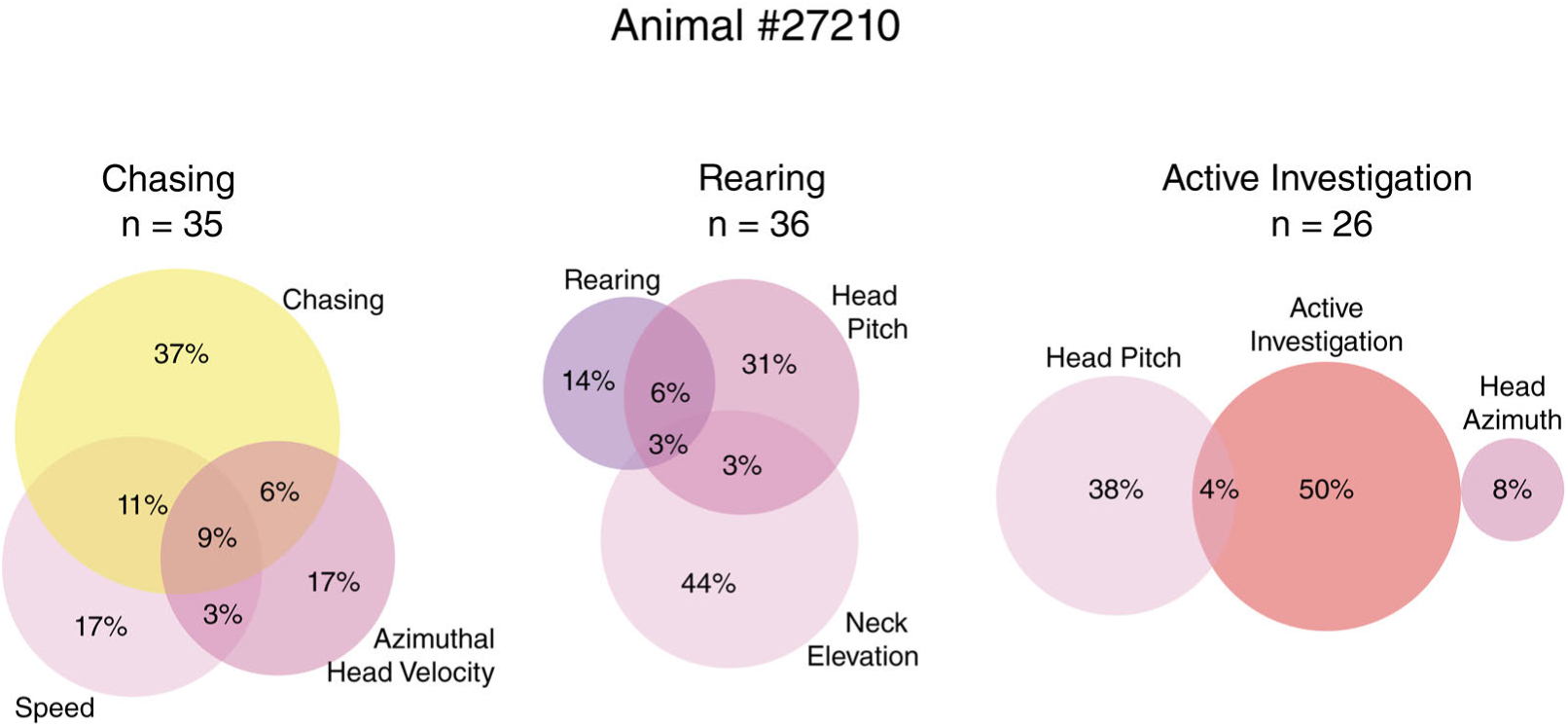
Overlap between units tuned to specific behavior states versus elementary kinematic features for animal #27210. Same as for Figures S12 and S13, but for the 3^rd^ animal in the study.

**Figure S15.**
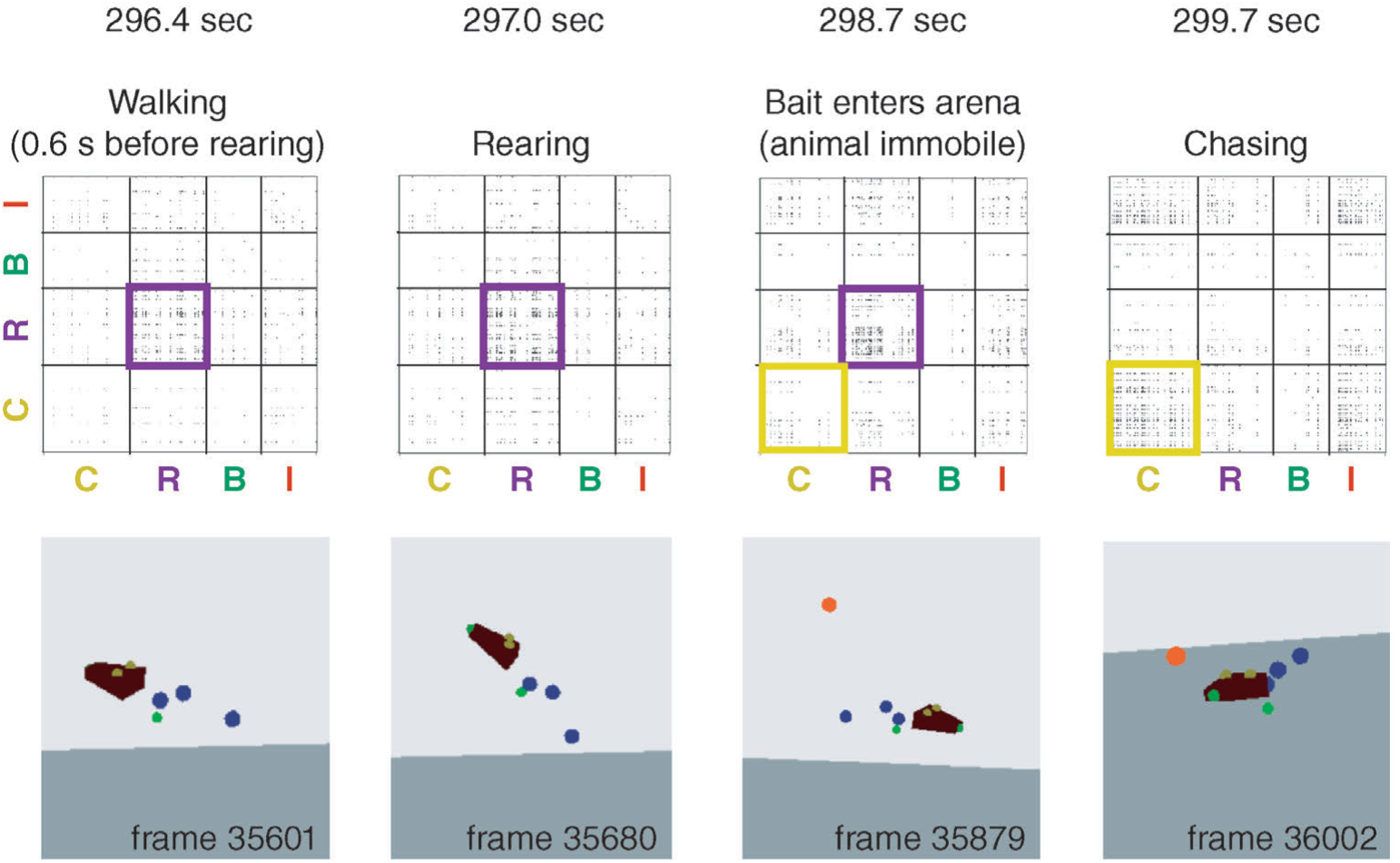
Behavior-specific mPFC coactivity patterns precede the onset of behaviors and transition smoothly between them. (Left) The ”rearing” coactivity matrix is expressed *>*0.5 s in advance of rearing, while the animal is walking on all fours, and (second from left) stays active when the animal follows through with rearing. (Second from right) Neural coactivity transitions from the ”rearing” matrix to the ”chasing” matrix as the bait (marked as red dot) is lowered into the arena; the animal is back on all four feet and still immobile. (Right) The ”rearing” graph dominates once the animal initiates pursuit of the bait.

**Figure S16.**
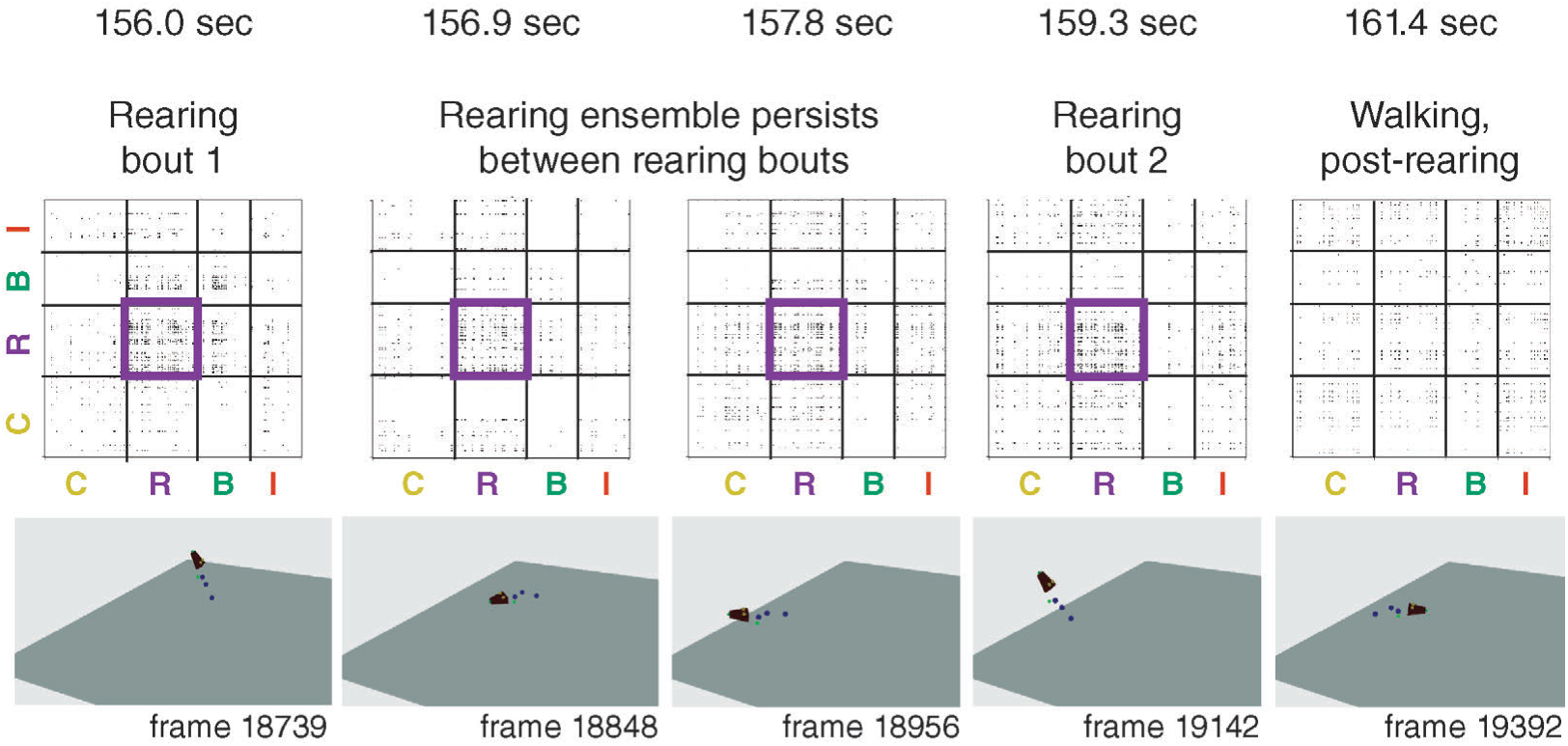
”Rearing” coactivity patterns persist between successive bouts of rearing. (Left panel) The neural ensemble for rearing is expressed during the first instance of a rearing bout, and (next two panels) is maintained while the animal walks on all fours before rearing a 2nd time (next panel). (Right panel) The rearing ensemble disappears after the two rearing bouts are over and the rat not is preparing to rear again.

**Figure S17.**
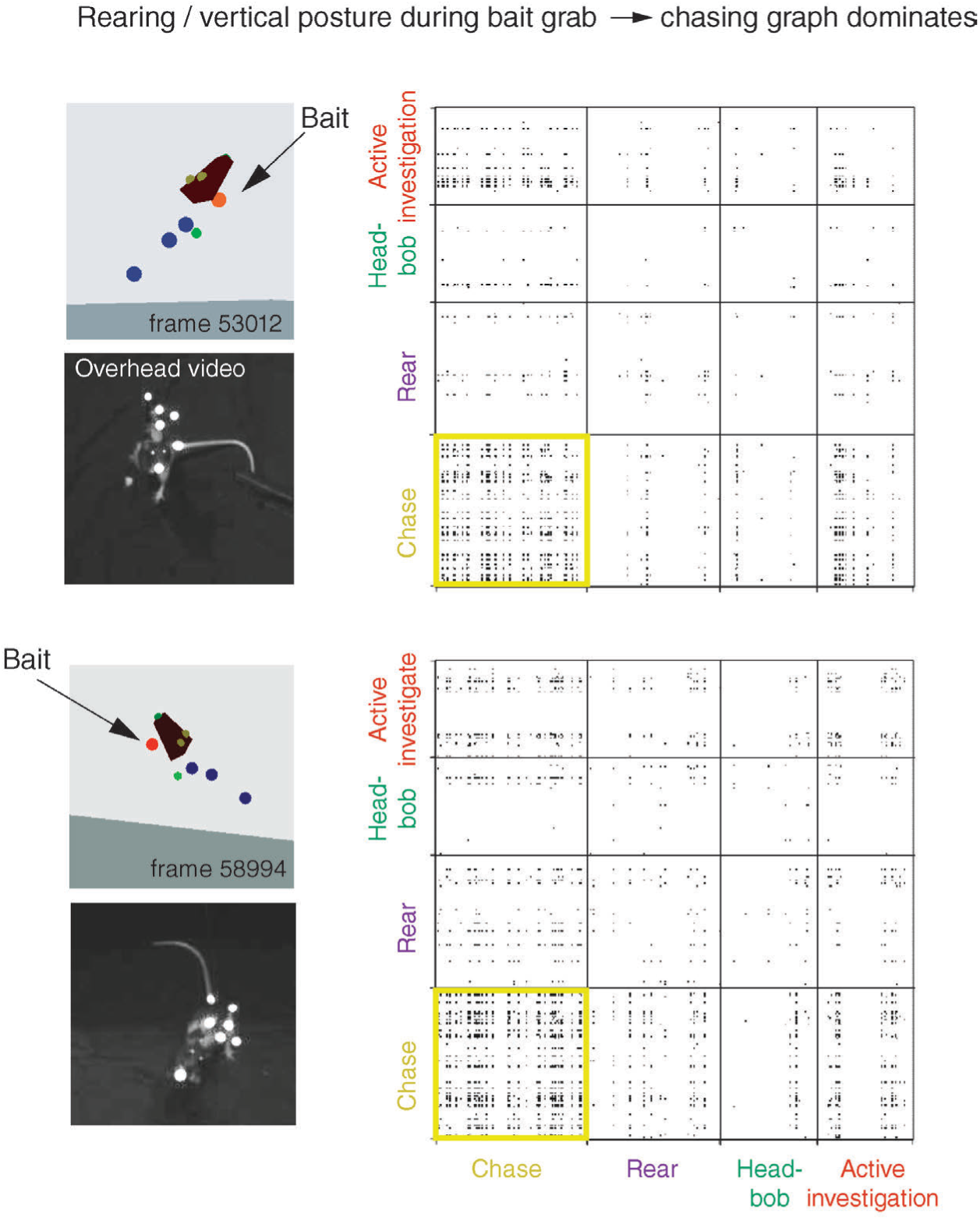
Neural coactivity patterns signify ”chasing” when animals rear to grab bait during chasing. Two examples from a chasing session where the animal reared (top) or raised its head (bottom) to grab the bait at then end of a chasing bout. Despite the vertical posture features, the population activity reflected ”chasing” rather than ”rearing”.

**Figure S18.**
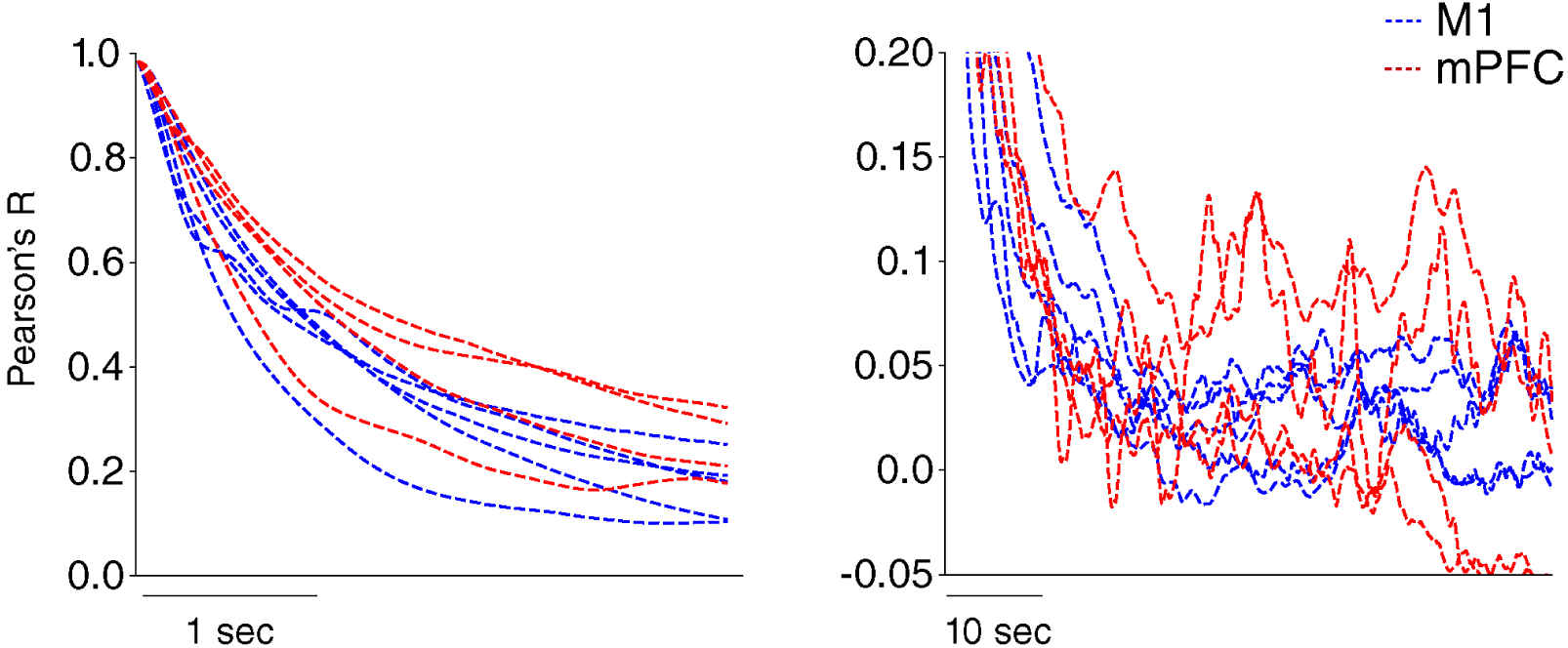
Temporal autocorrelations for kinematic features in M1 and mPFC data sets. Blue and red dashed lines show the post rescaling autocorrelation values for postural features (head pitch and azimuth; back pitch and azimuth) and running speed in the M1 (from Mimica et al., 2023) and mPFC recordings from open field sessions. Recordings in M1 were slowed by a factor of 0.5 in order to match kinematic profiles of the mPFC data set as shown. A 4 s autocorrelation window is shown (left), and the same is shown over 60 s (right).

**Figure S19.**
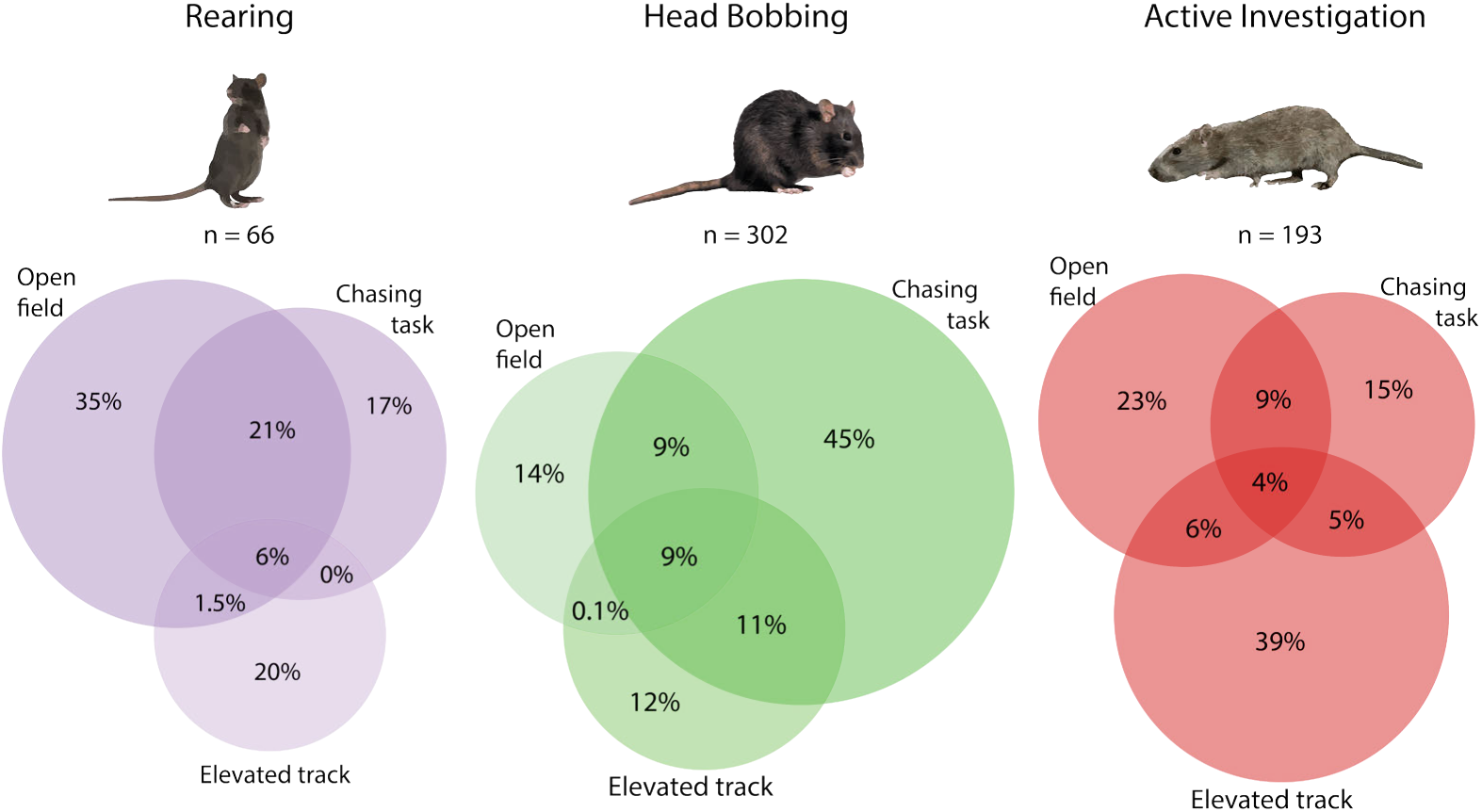
Behavioral selectivity of inhibited units also varied depending on the task in which the behavior was expressed. (left) Venn diagrams showing the overlap of neurons stably suppressed during rearing in each task, pooled across animals. (middle) Same for head bobbing, and (right) same for active investigation. The behaviors “rearing” and “active investigation” had a larger degree of overlap across the open field and chasing tasks; “head bobbing” had the largest overlap across the elevated track and chasing task.

**Figure S20.**
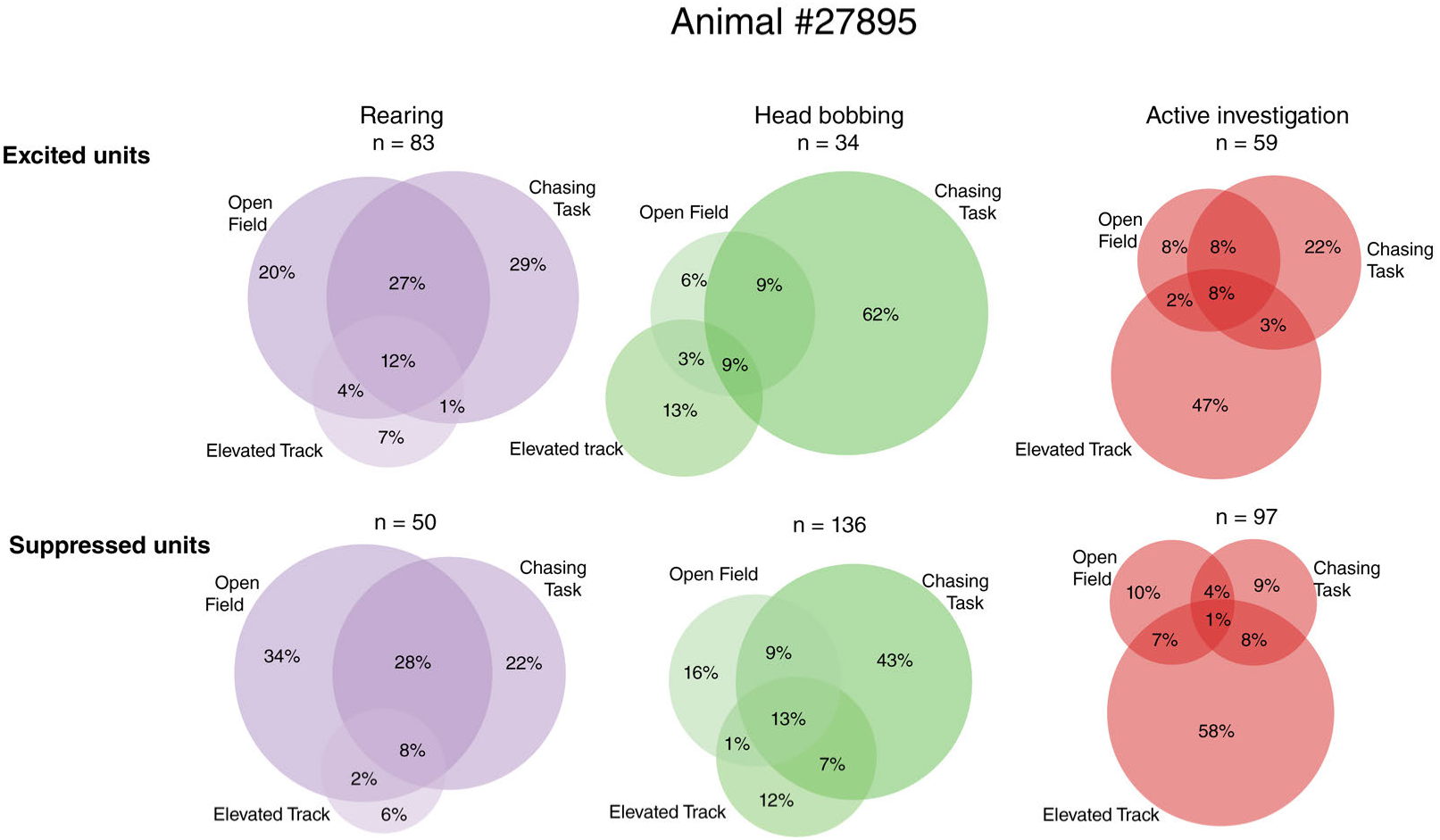
Venn diagrams showing across-task overlap of behaviorally-tuned neurons for animal #27895. (top row, left) Rearing-excited, (top row, middle) head bobbing-excited, (top row, right) active investigation-excited neurons. (Bottom row) same ordering as top row, but for behaviorally-suppressed neurons.

**Figure S21.**
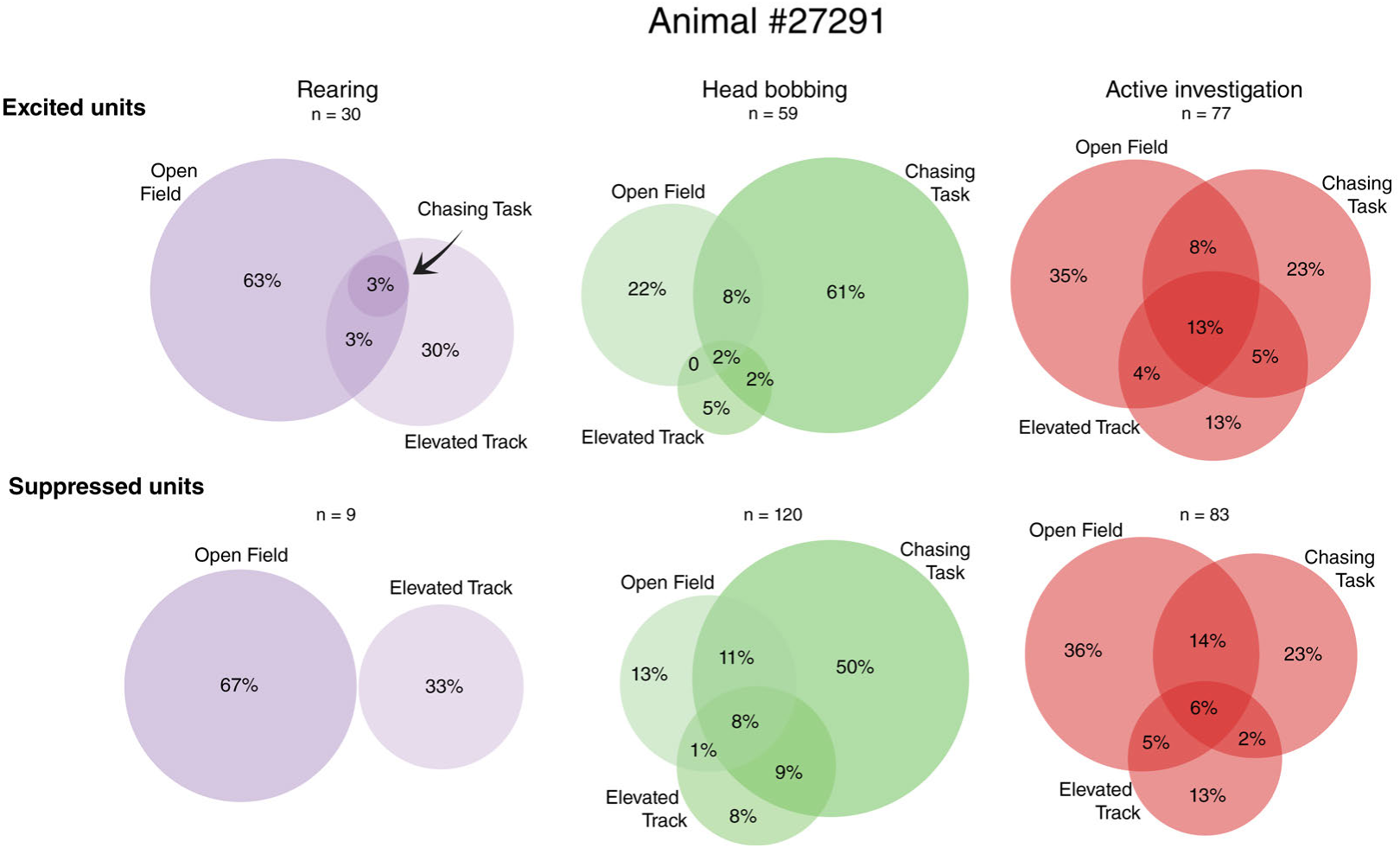
Venn diagrams showing across-task overlap of behaviorally-tuned neurons for animal #27291. (top row, left) Rearing-excited, (top row, middle) head bobbing-excited, (top row, right) active investigation-excited neurons. (Bottom row) same ordering as top row, but for behaviorally-suppressed neurons.

**Figure S22.**
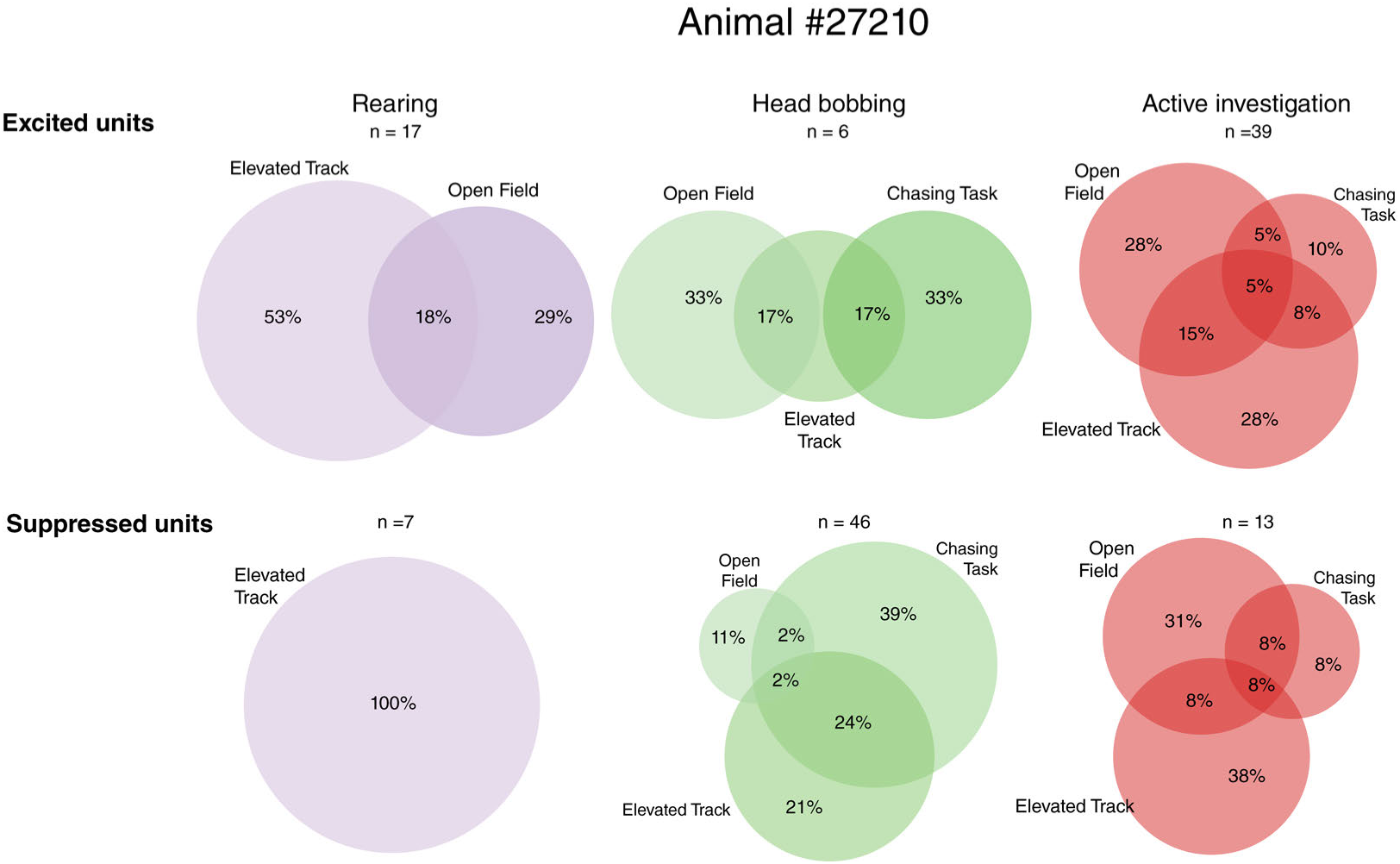
Venn diagrams showing across-task overlap of behaviorally-tuned neurons for animal #27210. (top row, left) Rearing-excited, (top row, middle) head bobbing-excited, (top row, right) active investigation-excited neurons. (Bottom row) same ordering as top row, but for behaviorally-suppressed neurons.

**Figure S23.**
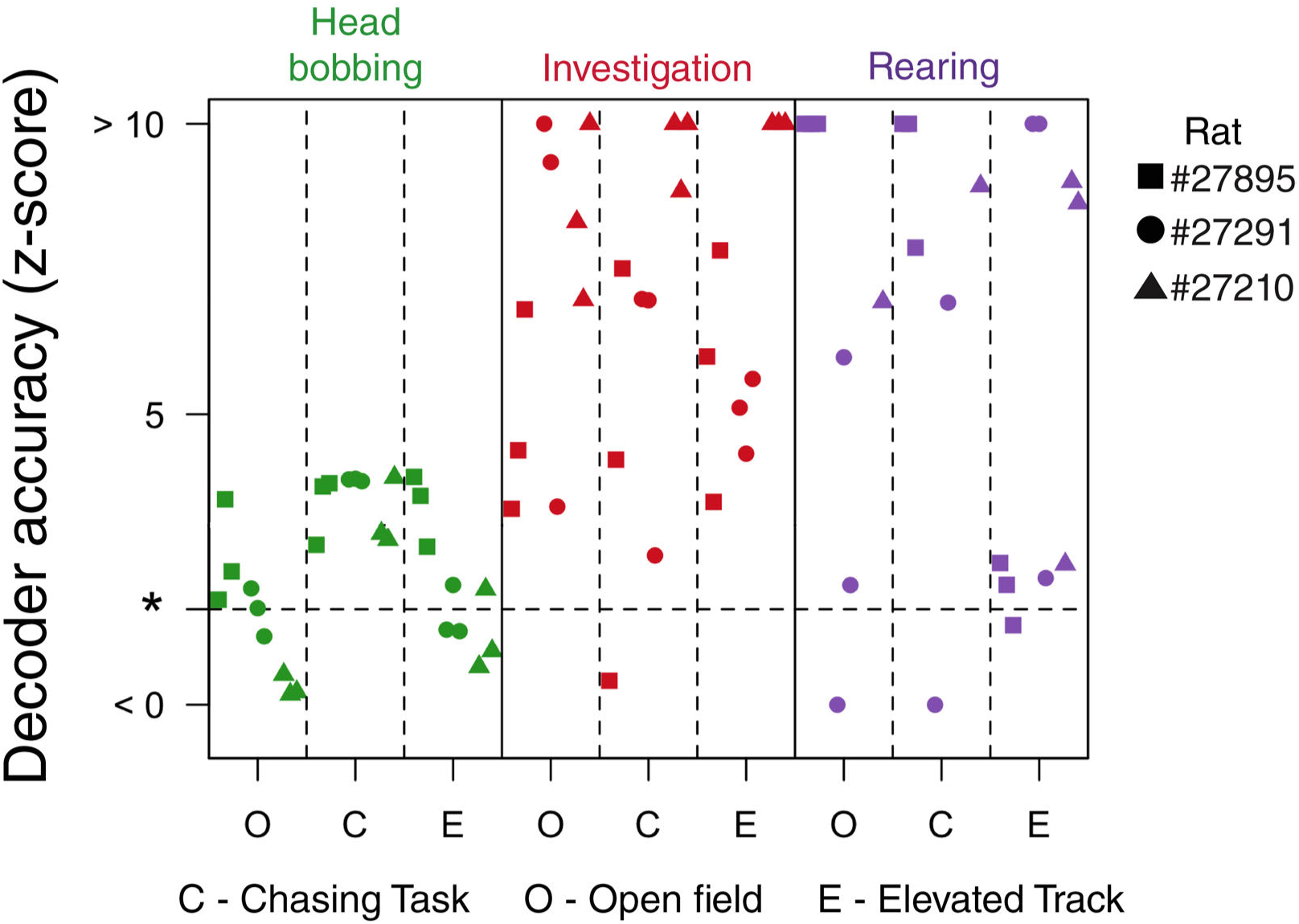
Scatter plot showing across-task z-scored decoding accuracy for each behavior from all animals, in all sessions. Each column is a session type, colors indicate specific behavior states, symbols denote animals. Dashed line indicates significance level of *α* = 0.05 for decoder performance.

## Notes

https://figshare.com/s/f09f8ac2cf9fff1f8f35

https://github.com/fredrine/behavior_state_mPFC_scripts

### Competing Interest Statement

The authors have declared no competing interest.

